# Activity during the first days of life predicts lifespan in a naturally clonal vertebrate

**DOI:** 10.64898/2026.02.21.707185

**Authors:** Ulrike Scherer, Sean M. Ehlman, David Bierbach, Jens Krause, Max Wolf

**Author notes:** Corresponding author Materials & Correspondence Correspondence and material requests may be addressed to Ulrike Scherer.

## Abstract

Lifespan varies widely among individuals, yet the extent to which such variation persists when genetic and environmental differences are minimized remains unclear. Here we quantify such stochastic lifespan variation in a naturally clonal vertebrate and test whether and how this variation is linked to early-life behavioral individuality. We followed *N* = 33 genetically identical Amazon mollies (*Poecilia formosa*), separated on day 1 of their life into highly standardized environments, from birth to death. Despite genetic uniformity and environmental standardization, lifespan varies markedly, spanning 502 – 826 days. Continuous high-resolution behavioral tracking during the first four weeks of life reveals that seemingly stochastic early-life activity differences explain 32.5% of this variation. Higher activity predicts shorter lifespan during the first two weeks, but as activity levels and among-individual variation in activity decline over early development, a U-shaped relationship emerges, with both low- and high-activity individuals outliving those with intermediate activity. These findings show that signatures of lifespan emerge within days of birth, even among genetically identical individuals, highlighting developmental stochasticity and early-life contingencies as major contributors to variation in life-history outcomes.

## MAIN

A central challenge in longevity and aging research lies in understanding why individuals differ so remarkedly in how long they live. Indeed, a formidable body of literature maps both the environmental and genetic correlates of lifespan variation in humans and other animals. Studies on human centenarians living in so-called ‘Blue Zones’ across the globe, for example, point to socio-environmental factors such as diet, lifestyle, and social structure that are associated with exceptionally long lifespan ^1^, while comparative and functional genomic work has identified specific genetic variants and their downstream metabolic pathways as a source of longevity variation both within and among species ^2–4^. Beyond their independent effects, genes and the environment have been shown to interact in powerful ways to shape lifespan ^5–7^. More recently, however, experimental work has suggested that traditional genetic and environmental explanations alone are insufficient to fully account for variation in lifespan among individuals. Using isogenic *Escherichia coli* and *Caenorhabditis elegans*, for example, it has been demonstrated that, even when eliminating genetic and environmental sources of variation as much as experimentally possible, substantial lifespan variation persists ^8–11^. These findings implicate stochastic processes (e.g., stochastic gene expression, developmental stochasticity), developmental contingencies, and ‘minor’ genetic and environmental/experiential differences as potential sources of lifespan variation among individuals. Up to now, however, virtually all of the experimental evidence comes from small and short-lived organisms like *E. coli* and *C. elegans*. Whether - and to what extent - similar pattern can be observed in longer-lived or even vertebrate animals is largely unknown.

In parallel, and largely independent from studies on stochastic variation in lifespan, recent work in the field of behavioral biology has documented a related phenomenon across a range of study systems: the emergence of “stochastic” behavioral individuality (consistent among-individual behavioral variation; ^12–15^) under conditions where genetic and environmental differences are reduced to a minimum ^16–26^. Genetically identical fish, for example, raised under near-identical environmental conditions, express consistent differences in activity and feeding behavior during the first weeks of life ^23–25^. Isogenic fruit flies, also reared in near-identical conditions, develop consistent individual differences in locomotor handedness, wing-folding, phototaxis, and object-fixated locomotion ^20–22^. Related results exist for inbred mice, clonal crayfish, clonal pea aphids, and *Daphnia spp.* ^16,27–31^. But while a substantial number of studies have characterized seemingly stochastic behavioral individuality, up to now, its significance and potential association with differences in long-term major life history outcomes such as lifespan remain virtually unexplored (but see Scherer et al., 2023).

Here, we are aiming to bridge the fields of stochastic behavioral individuality and stochastic lifespan variation by employing a unique, naturally clonal vertebrate system, the Amazon molly (*Poecilia formosa*). We closely monitored *N* = 33 genetically identical Amazon mollies, reared individually and under highly standardized experimental conditions from day 1 of their life until death, i.e., day 826 of the experiment (death of last individual). Using a custom-developed high-resolution tracking system, we characterized in depth early-life activity (average distance moved) and feeding behavior (time spent feeding per day), tracking individuals continuously for 10 hours per day over the first four weeks (i.e., 28 days) of life ^32,33^. At two phases during the experiment (day 38 to 280 and day 393 to 599), individuals were allowed to reproduce, and we tracked the complete reproductive output of individuals (i.e., the number and size of all offspring produced; see Scherer et al. (2023) for a detailed analysis of the reproductive profiles during the first reproductive phase). Shortly before the second reproductive phase, when individuals were approximately one year old, we characterized adult activity and feeding patterns. Throughout the experiment, we recorded body sizes of individuals in regular intervals. We recorded the day of death for each individual. We then asked: (1) to what extent do we observe differences in lifespan despite the lack of apparent genetic or environmental differences among individuals? And, (2) is stochastic early-life behavioral individuality in activity profiles or feeding behavior linked to lifespan variation?

## RESULTS

### Lifespan variation among genetically identical individuals

Our *N* = 33 genetically identical individuals, reared individually and under highly standardized environmental conditions, differ considerably in lifespan (**Figure 1**), ranging from 502 to 826 days (death of first and last individual, respectively; mean ± SD = 652 ± 79 days, coefficient of variation = 12.1 %). We identified 5 individuals as outliers (gray bars in **Figure 1**) using the ‘boxplot method’, which defines outliers by having their day of death beyond 1.5x the interquartile range. We excluded these outlier individuals from our main analyses as these individuals presumably died prematurely, i.e., for reasons other than physiological aging processes (original sample size including outliers: *N* = 38 individuals, lifespan range: 157 to 826 days; mean ± SD = 597 ± 165 days, coefficient of variation = 27.7 %).

**Figure 1.**
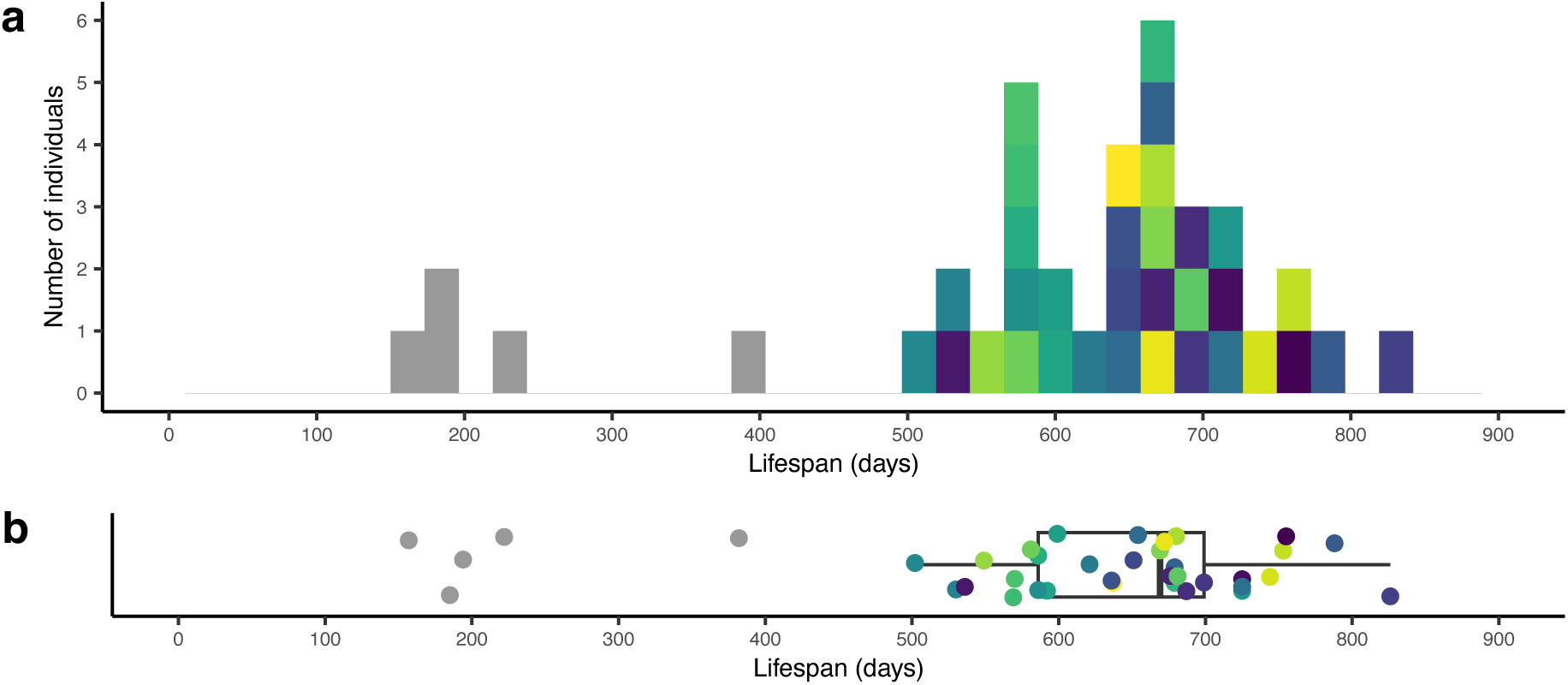
Variation in lifespan among genetically identical individuals raised individually and under near-identical conditions. (a) Histogram with the number of deaths over time for *N* = 38 naturally clonal Amazon mollies separated directly after birth into highly standardized individual environments; bars represent the number of individuals per bin. (b) The integrated boxplot displays median lifespan (middle line), 25th to 75th percentile (box), and 5th to 95th percentile (whiskers) as well as the raw data (points) for each individual. (a, b) Bar and point color corresponds to fish identity (*N* = 33), consistent across panels and figures (except *N* = 5 outliers in gray).

### Activity during the first days of life predicts lifespan – but no effect of feeding

As reported previously ^24^, even though individuals are genetically identical and were raised individually under highly standardized conditions from day 1 of their life onwards, we observe consistent among-individual differences in both activity and feeding behavior expressed over the first 28 days (i.e., four weeks) of life (activity: raw repeatability = 0.361, 95% CI = [0.316, 0.403], repeatability adjusted for age and body size = 0.491, 95% CI = [0.444, 0.547]; feeding: raw repeatability = 0.186, 95% CI = [0.147, 0.228], repeatability adjusted for age and body size = 0.249, 95% CI = [0.201, 0.301]) (**Supplementary Figure 1a, c**).

Among-individual variation in activity expressed over the first four weeks of life explains 32.5% (*R*^2^) of the variation in lifespan, with both high- and low-activity individuals living longer than those with intermediate levels of activity (*p* = 0.006, **Figure 2a**, **Supplementary Table 1**). To further assess the temporal development of this link, we next analyzed each week separately. We find that activity in week 1, week 2, and week 3 explains 28.0%, 15.0%, and 18.6% of the observed lifespan variation (*R*^2^), respectively, whereas activity in week 4 is not predictive of lifespan (**Figure 2b**; **Supplementary Table 1**). This week-by-week analysis reveals two key patterns. First, activity levels decline over time, resulting in a reduction of among-individual variation (**Figure 2b**, **Supplementary Figure 1**). Second, the shape of the activity– lifespan relationship shifts across development: during the first two weeks of life, activity is monotonically linked to lifespan, with more active fish dying sooner, whereas in week 3 - when behavioral variation is lowest - the relationship becomes non-monotonic, such that both low- and high-activity individuals outlive those with intermediate activity levels (**Figure 2b**; **Supplementary Table 1**). Notably, the same patterns emerge at a finer temporal scale: day-by-day analyses reveal a robust negative activity-lifespan relationships during the first days of life (days 2-10, except day 4), with quadratic relationships appearing mostly on later days (days 4, 17, and 20), when among-individual variation in activity is reduced (**Supplementary Figure 3, Supplementary Table 3**).

**Figure 2.**
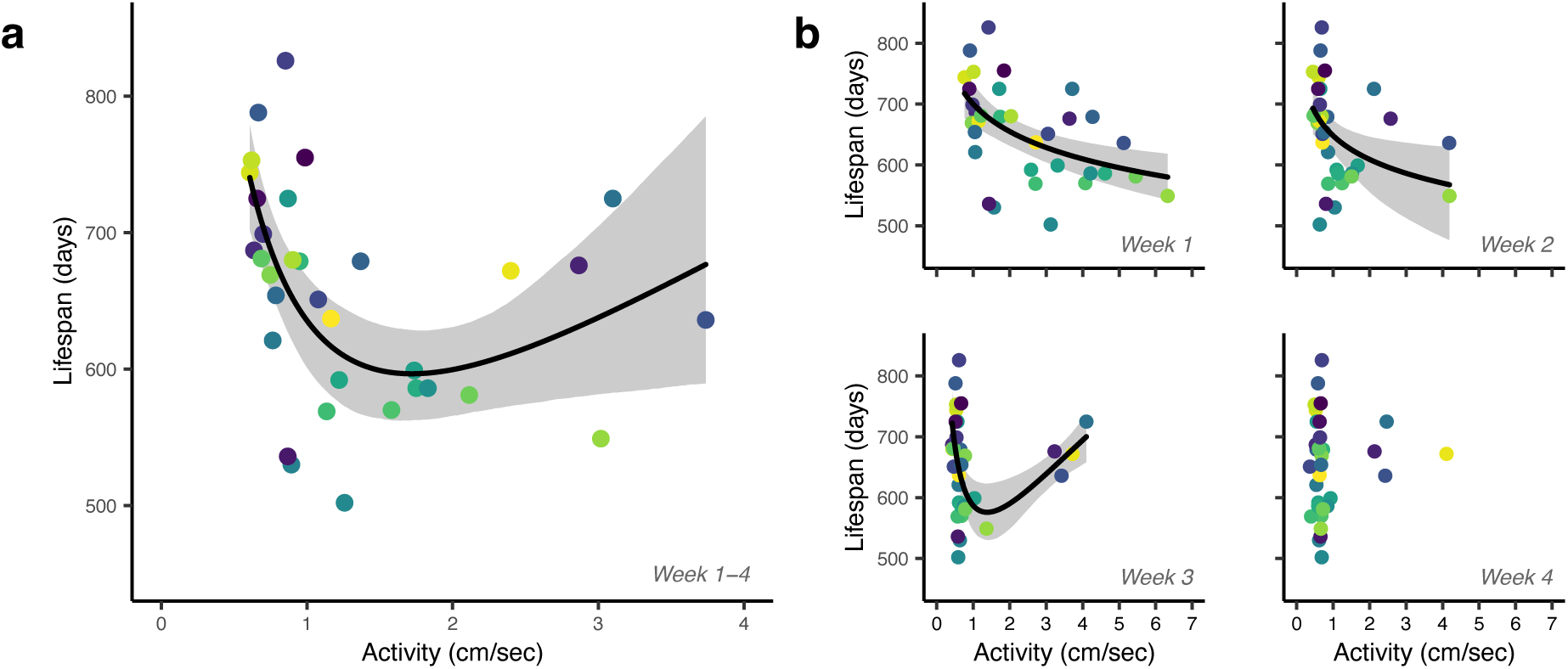
Early-life activity predicts lifespan. (a) Average activity expressed over the first four weeks of life explains 32.5% (*R²*) of the observed variation in lifespan, with individuals exhibiting either low or high activity living longer than those with intermediate activity levels. (b) At weekly resolution, the shape of this relationship shifts from negative (week 1-2) to quadratic (week 3), before becoming no longer detectable (week 4). (a, b) X-axes differ in scale. Regression lines (black) with 95% confidence intervals (gray) are shown only for significant relationships. Points represent raw data. Activity values were log-transformed for statistical analyses and back-transformed for plotting to aid interpretability. Data point color corresponds to fish identity, consistent across panels and figures (*N* = 33 individuals throughout).

Time spent feeding shows no direct association with lifespan, whether assessed as a four-week average or at weekly resolution (**Figure 3a-b**, **Supplementary Table 1**). However, we detect an indirect feeding-lifespan link, mediated via growth: fish that spend more time feeding over the first four weeks of life, grow to a larger maximum body size (*p* = 0.035, *R^2^* = 0.112**, Supplementary Table 4**), and fish that grow larger live longer (*p* = 0.020, partial *R^2^* = 0.106, **Supplementary Table 5**). In contrast to the substantial effect of early-life activity on lifespan, explaining 32.5% of the observed variation in lifespan (see above), the indirect feeding-lifespan link explains only about 1% of the variation in lifespan. A week-by-week analysis reveals that the link between feeding behavior and body size is strongest in week 3 of life, while feeding behavior in weeks 1 and 4 exhibits non-significant trends, and no association is detected in week 1 (**Supplementary Note 4**). Maximum body size was estimated from individual van Bertalanffy growth models. Importantly, size measurements were curtailed to the last measurement when all individuals were still alive, thereby standardizing the growth period over all individuals (for more details see Methods).

**Figure 3.**
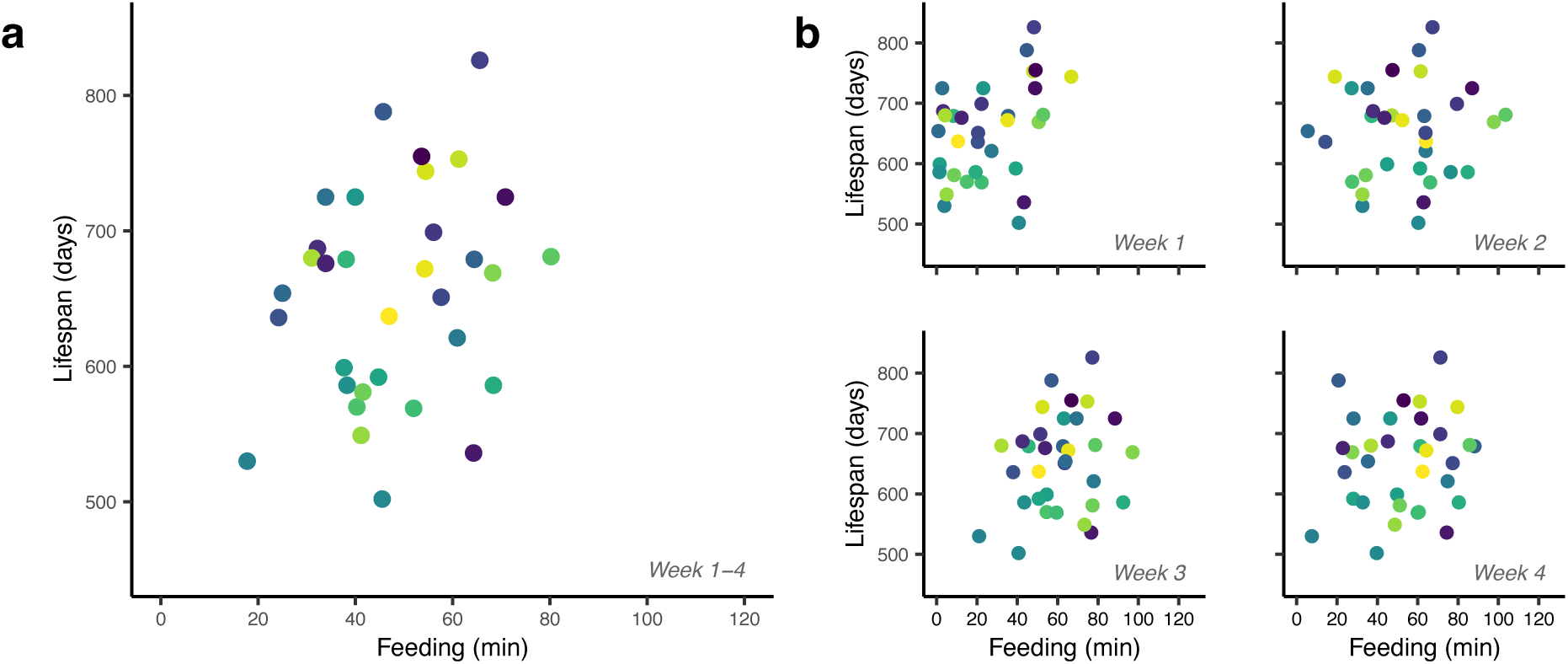
Early-life feeding behavior does not predict lifespan. (a) Average time spent feeding per day does not predict lifespan – whether expressed over (a) the first four weeks of life or at (b) weekly resolution. (a, b) Points represent raw data (*N* = 33 individuals throughout). Data point color corresponds to fish identity, consistent across panels and figures.

All our results above are robust to controlling for potential variation in size at birth. Specifically, when including size at birth in our models above, the key patterns remain unchanged: the activity–lifespan relationship remains initially negative before shifting to a U-shape, and feeding behavior continues to show no direct association with lifespan (**Supplementary Table 6**). Size at birth was estimated from individual van Bertalanffy growth models (see Methods).

As noted above, we excluded five outlier individuals that likely died for reasons other than physiological aging (**Figure 1**). When including these outliers in our analyses, we continue to find a U-shaped link between activity expressed over the first four weeks of life and lifespan, whereas no activity-lifespan link is detected when restricting our data to single weeks (**Supplementary Table 7**). Furthermore, when outliers are included, we observe a positive association between feeding behavior and lifespan in the first week of life, but this link is absent when considering the full four-week period or any other week (**Supplementary Table 7**).

We do not detect any effects of long-term housing tanks on lifespan (**Supplementary Table 2**). All our results above are controlled for descent (i.e., maternal identity) (**Supplementary Tables 1-7**).

### Beyond early-life behavior: lifespan in relation to additional behavioral and reproductive measures

While the present study focuses on the link between early-life behavior and lifespan, individuals used in this study were also characterized for reproductive output ^24^ and several additional behavioral measures, in accordance with 3R guidelines that maximize data collected from individual animals. Additional behavioral measures include a 9-day period of repeated standard behavioral assays, performed immediately following early-life behavior tracking, as well as later-in-life activity and feeding tracking (see **Supplementary Table 3** for a detailed overview of all experimental phases). All phenotyping was conducted in a highly standardized manner across all individuals. For completeness, we here report analyses assessing potential links among these additional traits and early-life behavior and/or lifespan. These analyses are intended as a high-level overview, accordingly, we focus on behavior averaged across the first four weeks of life only and do not consider weekly or daily variation for these additional analyses.

Early life behavior (activity and feeding) does not predict behavior expressed during standard behavioral assays (activity in a new tank, novel object response, and sociability), except for a positive correlation between early-life activity and activity in a new tank (**Supplementary Table 9**). We find no link between behavior expressed during these behavioral assays (activity in a new tank, novel object response, and sociability) and lifespan (**Supplementary Table 10**).

Later-in-life behavior is not correlated with early-life behavior (activity and feeding, respectively) (**Supplementary Tables 11**). Nevertheless, later-in-life activity exhibits a U-shaped relationship with lifespan, with both low- and high-activity individuals living longer than those with intermediate activity levels - mirroring the pattern observed for early-life activity (**Supplementary Table 12**). Later-in-life feeding behavior is not associated with lifespan (**Supplementary Table 12**).

Reproductive characteristics (i.e., age of first reproduction, average brood size and average offspring size over all broods produced across life) are not related to lifespan, and we find no direct link between early-life differences in activity and feeding behavior and any of our three reproductive characteristics (**Supplementary Tables 13-14)** (analyzed in detail in Scherer et al. (2023) for the first reproductive phase).

We note that, for logistic reasons, the exact timing of later-in-life behavioral typing (Phase 4) and the second reproductive phase (Phase 5) varied among individuals from different mothers (see **Supplementary Table 3** for a detailed overview of all experimental phases with exact time windows), however, all analyses are controlled for maternal identity and this factor did not explain any variation in lifespan (**Supplementary Tables 1-2, 4-14**).

## DISCUSSION

We conducted a highly standardized, whole-life laboratory study, following genetically identical Amazon mollies - separated directly after birth into near-identical environments – from their first day of life until their death more than two years later. Remarkably, despite the lack of detectable genetic and environmental differences among these individuals, (1) we find substantial variation in lifespan. Moreover, (2) we find that seemingly stochastic differences in early-life activity patterns predict a substantial proportion of the observed differences in lifespan (roughly 33%), with the shape of the relationship shifting from negative-monotonic to quadratic over early development – coinciding with declining among-individual variation in activity. Feeding behavior, in contrast, had only a very minor indirect effect on lifespan, mediated via growth, explaining roughly 1% of the lifespan variation.

We find that our naturally clonal fish, raised in near-identical environments differ substantially in their lifespan. While previous findings in the bacteria *E. coli* and the nematode *C. elegans*, have already documented such seemingly stochastic lifespan differences (i.e., differences in lifespan that cannot be attributed to genetic or environmental differences and their underlying cellular, molecular, or developmental mechanisms ^8–11^, our study significantly advances these previous findings by showing that stochastic lifespan variation can also arise in a vertebrate system, thus demonstrating that such variation is not restricted to the narrow set of previously studied taxa or a specific rearing paradigm and that stochasticity may play a fundamental role in shaping lifespan variation across a broader range of species.

Our study demonstrates that stochastic variation in lifespan is foreshadowed by activity expressed as early as the very first days of life. Two aspects of this finding are particularly compelling: (i) the magnitude and (ii) robustness of this link. (i) Early-life activity during the first four weeks of life explains roughly one third of the observed variation in lifespan – given that this behavioral variation is seemingly stochastic (i.e., no apparent genetic and environmental variation among individuals), the observed level of explanatory power is remarkable, and it suggests that early-life activity reflects more fundamental developmental or physiological differences that become established very early and shape long-term trajectories. Because any such differences must arise from processes other than genetic or major environmental variation, our finding heavily underscores the potency of processes like stochastic gene expression, developmental stochasticity, and minor environmental/experiential differences, in generating variation in key life-history outcomes. (ii) Intriguingly, the activity-lifespan relationship proves remarkably robust across temporal scales: the pattern observed at a weekly resolution (i.e., a negative-monotonic link that becomes U-shaped and then not detectable anymore) remains detectable even when the data are partitioned into single days – with a single day (10 hours per individual) constituting only a small fraction of the data used to assess weekly averages (70 hours per individual). Yet, it still carries predictive information about lifespan measured years later. The fact that such a strong signal emerges even from minimal behavioral sampling further underscores that early-life activity is not merely a noisy snapshot but reflects stable differences in underlying developmental or physiological pathways that are established very early in life.

Beyond the strength and robustness of the observed activity–lifespan link, we found its shape to be developmentally dynamic. During the first two weeks of life, higher activity is negatively associated with lifespan, with more active individuals dying sooner. This finding is in line with predictions from the pace-of-life-syndrome (POLS) framework, which proposes that individuals fall along a continuum from fast to slow “paces of life”; in contrast to standard pace-of-life predictions, however, we neither find a direct link between behavior and reproductive characteristics nor a link between reproductive characteristics and lifespan – thereby adding to the broadly mixed evidence for pace-of-life predictions ^34–37^. As development proceeds, the relationship between activity and lifespan shifts from negative monotonic to a U-shaped link, coinciding with a marked reduction in behavioral variation: most individuals converge on low activity levels, while only a small subset of individuals remains more active. Intriguingly, these active individuals live longer than expected under a simple negative-monotonic model. Given that high or low average trait values are typically associated with greater behavioral consistency than intermediate ones, a possible interpretation is that individuals that more consistently exhibited low- and high activity levels benefitted from higher behavioral consistency, e.g., through reduced physiological costs associated with behavioral switching ^38,39^. Rather than reflecting a fixed activity-lifespan trade-off, our findings suggest that the shape of this relationship may be context-dependent. In particular, convergence toward similar behavioral phenotypes - and the emergence of nonlinear associations - may not occur under all environmental conditions. Disentangling how environmental context shapes both stochastic behavioral variation and its long-term consequences for lifespan will be a critical task for future experimental work.

We found no direct link between time spent feeding and lifespan. However, there was a subtle, indirect link: individuals that consistently spent more time feeding during the first four weeks of life grew to a larger final size (over a standardized growth period that was identical for all fish), and larger fish lived longer. This suggests that higher food intake may enhance body condition, which in turn supports longevity. Yet, this link between feeding behavior and lifespan, mediated via body size, was quite weak, explaining only about 1% of the variation in lifespan. The weakness of this effect likely reflects our experimental design, in which food was abundantly available at a fixed location, thereby minimizing variation in food acquisition and eliminating the need for active food search. In more natural environments, where food is patchy, scarce, or unpredictable, differences in foraging behavior are likely to translate into greater variation in energy intake and body condition, potentially strengthening links between feeding-related behaviors and lifespan. This highlights the importance of considering ecologically realistic environments when investigating how stochastic behavioral differences influence life-history outcomes in future studies.

Taken together, our findings reveal that (i) even under the most standardised conditions, where genetic and environmental differences are minimized, individuals differ markedly in their lifespan; and that (ii) signatures of this major long-term life outcome are detectable within the first weeks of life. The early-life activity-lifespan link is both surprisingly strong and robust. A shift from negative-monotonic to a U-shape, coinciding with declining variation in activity, illustrates the potential context-dependency of its shape. As the observed patterns emerge despite the absence of apparent genetic or environmental differences, our findings strongly suggests that pre-birth processes (stochastic gene expression, developmental stochasticity) and/or micro-environmental differences during the earliest stages of life can initiate developmental pathways that cascade into lifelong phenotypic differences. More broadly, our work adds to the growing recognition of stochasticity being not minor noise around genetically and environmentally determined variation, but a potent source of phenotypic variation itself. Understanding these early stochastic processes, and how they are amplified, canalized, or buffered across development, represents an important frontier for developmental and evolutionary biology, behavioral ecology, and the study of longevity and ageing.

## METHODS

All animal care and experimental protocols complied with local and federal laws and guidelines and were approved by the appropriate governing body in Berlin, Germany, the Landesamt fur Gesundheit und Soziales (LaGeSo G-0224/20).

### Study species and holding conditions

The Amazon molly is a naturally clonal freshwater fish that arose from a single hybridization event between the Atlantic molly, *Poecilia mexicana*, and the sailfin molly, *Poecilia latipinna*, roughly 120,000–280,000 years ago ^40–42^. As a gynogenetic species, the Amazon molly relies on sperm from closely related, sexually reproducing species to trigger embryonic development, though the male’s sperm is not incorporated into the offspring’s genome ^41,43–46^.

Test individuals were obtained from an Amazon molly stock at Humboldt-Universität zu Berlin, (Berlin, Germany). The lineage used in our experiment was established 6 months prior to the experiment, by housing a single Amazon molly from the stock population along with a *Poecilia mexicana* male and all resulting offspring in a 50-liter tank (∼20–50 fish). To initiate the experiment, potential mothers were separated from this isolated lineage to give birth in individual tanks. This separation enabled the tracking of mother identity (*N* = 3) for individuals utilized in our experiment. Throughout, i.e., in the husbandry as well as during the experiment, we maintained standardized conditions with a 12:12h light:dark cycle, air temperature control (approx. 24±1°C), and weekly water changes. In the husbandry, fish were fed ad libitum twice per day on five days per week with powder food (Sera vipan baby).

### General overview

We employed a whole-life experimental design, following *N* = 33 genetically identical individuals from birth until death under highly standardized laboratory conditions. While the present study focuses on the link between early-life behavior and lifespan, the data that were analysed here were collected as a part of a broader phenotyping project, aimed at quantifying multiple behavioral and life-history traits across development (Scherer et al. (2023), manuscripts in preparation). Thus, across the full project, individuals experienced six highly standardized experimental phases (see **Supplementary Table 3** for a detailed overview of all experimental phases with exact time windows). Only Phase 1 (early-life behavioral typing) and Phase 6 (lifespan monitoring) are central to this study; the remaining phases (including standardized behavioral assays, later-in-life behavioral typing, and two reproductive periods) are described briefly below for completeness.

Except for brief periods of behavioral typing (Phases 1, 2, and 4; see below), individuals were maintained in individual housing tanks (11 liter), visually separated yet connected via a single flow-through water system. Each housing tank was equipped with a small plastic pipe (4 x 2 cm) and a bundle of ‘Sera Biofibres’, which are structurally equivalent to thread algae. Within the tank system, individual tank locations were either central (with a neighbouring tank on both sides) or peripheral (a neighbouring tank on one side only), and tanks were dispersed over 3 vertical levels (top, middle, and bottom level). Tank centrality and level affected neither reproductive characteristics ^24^ nor lifespan (**Supplementary Table 2**). In those individual housing tanks, fish were fed twice a day for five days per week with powder food (Sera vipan baby). Per feeding, each individual received 1/64 tsp (up to the age of 70 days) or 1/32 tsp (from the age of 70 onwards until death) of food.

The original dataset comprised *N* = 45 individuals. For all analyses, we removed *N* = 6 individuals who died shortly after the transfer to individual housing and breeding tanks (Phase 2), when they were <44 days old; as well as *N* = 1 female that did not die a natural death (age 112 days). Of the remaining 38 individuals, we identified five outliers, all exhibiting lifespans shorter than 1.5x the interquartile range of the population distribution of lifespans (**Figure 1**). Our final analysis thus focused on *N* = 33 individuals (for a robustness analysis with respect to the exclusion of outliers see ‘Results’ or **Supplementary Note 6**).

### Experimental phases

#### Early-life behavioral typing (Phase 1)

We quantified individual activity and feeding behavior over the first 28 days of life (i.e., four weeks) using a high-resolution, long-term tracking system ^32,33^. Newborns were placed in separate, identical observation tanks on the day of birth (**Supplementary Figure 2**). Starting the next day (i.e., the first full day of life), activity was recorded over the first 8 hours of each day, followed by a 2-hour feeding recording (Basler acA5472 cameras with 5 frames per second). During the feeding period, individuals were presented with a ‘food patch’, that was positioned at a standardized location in the tank (**Supplementary Figure 2**). Food patches were prepared every 2 to 3 days using Sera vipan baby powder food and agar (protocol see Scherer et al. (2023)). Recordings were tracked using the software Biotracker (Mönck et al., 2018), and resulting tracking data (movement data in the form of csv-files with xy-coordinates over time) were processed (including visualisation of tracking data and calculation of metrics) using a custom repository developed for this purpose ^32^. Individual activity was assessed as average daily swimming speed (cm/sec), and the time spent feeding was determined as the duration individuals spent in the ‘feeding zone’ (minutes), i.e., a 5 x 13 cm large zone surrounding the food patch. Individuals were fed 7 days per week during this phase. Observation tanks were cleaned once a week, starting approx. 2 hours after the feeding. To do so, tanks were scrubbed with a soft sponge, all dirt was removed with a hose and tanks were refilled with aged, tempered tap water. For the duration of the cleaning, individuals were transferred to 2L-plastic boxes (cased with Armaflex insulation).

#### Behavioral Assays (Phase 2)

Over a period of nine days, all individuals completed a series of standard behavioral assays, being tested three times in each of three classic paradigms (totalling nine behavioral tests per individual, with one test per day): activity in a new, unfamiliar tank, response to a novel object (average distance towards a novel object that was introduced into the tank), and sociability (time spent near a small, clear container housing two similar-sized, unknown conspecifics introduced in the tank). All assays were conducted in the above-described observations tanks (see Phase 1). See **Supplementary note 9** for a detailed description of the procedure.

#### Reproduction I (Phase 3)

Individuals were allowed to reproduce over a period of approx. 8 months. As Amazon mollies need sperm from a closely related species to initiate reproduction (see ‘Study species and holding conditions’), each housing tank was endowed with one *P. mexicana* male. Females were swapped between housing tanks (and males) once a week allowing us to experimentally control for potential tank/male effects on female reproductive profiles (number and size of offspring produced over multiple broods) (described in detail in Scherer et al. (2023)).

#### Later-in-life behavioral typing (Phase 4)

We characterized adult behavior (activity, time spent feeding) for an additional 9 days as described in the above ‘Early-life behavioral typing’ (Phase 1). In order to do so, as a preparatory step, females were housed without males for approx. 3 months to ensure they were not gravid during the following behavioral recordings. Different to Phase 1, the size of the feeding zone (10 x 13 cm) was increased to match adult fish sizes. Phase 4 was initiated at slightly different points in time (a difference of two days) for individuals from Mom 1 and 2 (age = 369 days) vs. individuals from Mom 3 (age = 371 days). Importantly, all our analyses are controlled for maternal identity, and this factor did not explain any of the variation in lifespan (**Supplementary Tables 1-2, 4-15**).

#### Reproduction II (Phase 5)

Females remained in their respective housing tank while a single *P. mexicana* male was added for a period of approx. 5 months. We note that for logistic reasons, this differed from Reproduction I, where females were swapped among males/housing tanks once a week (see above). For individuals from Mom 1 and 2, this phase was initiated at age = 445 days, while Mom 3 individuals started at age = 393, however, all our analyses are controlled for maternal identity, and this factor did not explain any of the variation in lifespan (**Supplementary Tables 1-2, 4-15**). We note that we recorded complete reproductive profiles for all females across their entire lives, i.e., we also collected data (number and size of all offspring) on any broods produced outside the designated reproductive phases.

#### Individual housing (Phase 6)

Females were housed individually until death.

### Lifespan monitoring and body-size measurements

While we recorded the day of death for each individual as closely as possible to reflect natural lifespan, a small number of individuals (*N* = 8) had to be euthanized as mandated by animal welfare regulations to prevent pain and suffering (following procedures outlined in ^48^). Crucially, all eight of these cases were the result of severe, unrecoverable, end-of-life-associated illnesses that are expected to result in death within a few days if left untreated. Thus, euthanizing these individuals introduced only a minimal deviation in the recorded date of death. Specifically, *N* = 6 females were sacrificed due to severe dropsy (characterized by swelling of the body, loss of appetite and impaired ability to swim), *N* = 1 female was euthanized after displaying signs of extreme emaciation, coupled with loss of motor control, and *N* = 1 female was euthanized because of an intestinal prolapse and abnormal behavior. For *N* = 3 females, the exact day of death was not recorded. In these cases, we inferred the day of death from their sequential size measurements, assigning it to the midpoint between the last measurement confirming they were alive and the subsequent scheduled measurement in which they were absent. These consecutive size measurements were taken 28, 42, and 43 days apart, respectively, yielding a maximum uncertainty of 14-22 days in these three death date estimates. To confirm the robustness of our findings, we repeated the main analyses (testing for a link between early-life beavhior and lifespan) excluding these three individuals and found qualitatively the same results, i.e., an activity-lifespan link that shifts from negative-monotonic (week 1 and 2) to U-shaped (week 3 and over all four weeks) (**Supplementary Table 8**).

Throughout, individuals were measured for body size regularly (weekly size measurements during Phase 1-3, monthly size measurements during Phase 4-6) using ImageJ ^49^. Body size was measured as standard length, i.e., the distance from the snout to end of the caudal peduncle. We modelled individual growth curves employing the von Bertalanffy growth model ^50^, a widely utilized logistic function for fish growth. The parameters estimated within this function include the theoretical age when size is zero (*t_0_*), the growth coefficient (*K*), and the maximum predicted size. To estimate individual growth curves, we curtailed size measurements to the last measurement when all individuals were still alive (day 493, mean ± SD number of measurements per individual = 20 ± 6).

### Statistical analyses

Data analyses were performed in R version 4.2.1 ^51^ using the *lme4*-package for modelling ^52^, *sjPlot* for creating model summary tables ^53^, *ggplot2* for plotting ^54^, and *partR2* for calculating partial *R^2^* values ^55^. For all analyses, activity was log-transformed (natural logarithm) for normality. Model fit for all linear-mixed-effect models was assured via inspection of residual and qq-plots. Most parsimonious models were fit via stepwise-backward removal of non-significant predictors. In the main text, we report p-values for significant predictors only. Complete model summary tables including estimates, test statistics, and p-values for all predictors (for full and final, i.e., most parsimonious models) are presented in the supplement (**Supplementary Tables 1-2, 4-15**).

Behavioral repeatabilities were estimated following Hertel et al. (2020). We calculated raw repeatability for early life behavior by building a linear mixed-effects model with the behavior of interest as response variable (daily log-activity and daily feeding behavior), and test fish identity as random term (*N* = 913 observation days from *N* = 33 individuals; i.e., for each behavior, *N* = 28 data points per individual, *N* = 11 missing values due to technical issues); no fixed effects were included. We then calculated adjusted repeatabilities by including age (week 1-4, categorical variable), body size (based on weekly size measurements), maternal identity, and a size-age interaction term (the effect of body size on behavior was dependent on age) as fixed effects in the above-described model.

We quantified variation in lifespan by calculating the coefficient of variation as: (standard deviation / mean) * 100. The coefficient of variation reflects the relative variability of lifespan in relation to its mean. To test whether early-life behavior predicts lifespan, we built a linear mixed-effects model with activity (averaged over log-transformed) and feeding behavior as predictor variables and lifespan as the response variable. Both activity and feeding behavior were averaged across the first four weeks of life, resulting in *N* = 33 data points (i.e., one data point per individual). Based on visual inspection of the data, which suggested a U-shaped relationship between activity and lifespan, we also included a quadratic term for log-activity. Maternal identity was included as a random effect. Furthermore, to investigate the temporal development of the activity-lifespan relationship, we repeated the above model for each of the four weeks of early life, i.e., we averaged activity within each respective week before fitting the model (*N* = 33 data points per model).

## Acknowledgements

We gratefully acknowledge funding from the Deutsche Forschungsgemeinschaft under Germany’s Excellence Strategy EXC 2002/1, ‘Science of Intelligence’ (project number 390523135) and from a DFG ‘Eigene Stelle’ grant to SME (project number 536703956). Members of the Wolf and Krause labs at the Leibniz Institute of Freshwater Ecology and Inland Fisheries, Humboldt University, and SCIoI Exzellenzcluster provided invaluable help with animal husbandry, experimental design and logistics, as well as coding and data management.

## Author contributions

J.K. and M.W. acquired funding. All authors conceived and designed the experiment. U.S. conducted the experiment. U.S., S.E., and M.W. outlined the data analysis. U.S. conducted the data analysis. U.S. and M.W. wrote the first manuscript draft. All authors commented on the manuscript and contributed to the final version.

## REFERENCES

1. Argentieri, M. A., Amin, N., Nevado-Holgado, A. J., Sproviero, W., Collister, J. A., Keestra, S. M., Kuilman, M. M., Ginos, B. N. R., Ghanbari, M., Doherty, A., Hunter, D. J., Alvergne, A., & van Duijn, C. M. (2025). Integrating the environmental and genetic architectures of aging and mortality. Nature Medicine, 31(3), 1016–1025. 10.1038/s41591-024-03483-9

2. Barde, W., Renner, J., Emery, B., Khanzada, S., Hu, X., Garthe, A., Rünker, A. E., Amin, H., & Kempermann, G. (2025). Beyond nature, nurture, and chance: Individual agency shapes divergent learning biographies and brain connectome. Science Advances, 11(2). 10.1126/sciadv.ads7297

3. Bates, D., Mächler, M., Bolker, B., & Walker, S. (2015). Fitting linear mixed-effects models using lme4. Journal of Statistical Software, 67(1). 10.18637/jss.v067.i01

4. Bell, A. M., Hankison, S. J., & Laskowski, K. L. (2009). The repeatability of behaviour: a meta-analysis. Animal Behaviour, 77, 771–783. 10.1016/j.anbehav.2008.12.022

5. Bierbach, D., Laskowski, K. L., & Wolf, M. (2017). Behavioural individuality in clonal fish arises despite near-identical rearing conditions. Nature Communications, 8(1), 15361. 10.1038/ncomms15361

6. Bierbach, D., Schutz, C., Scherer, U., & Pacher, K. (2024). Standards for scientific fish keeping: Poeciliid fishes with special focus on the guppy (*Poecilia reticulata*) and mollies ((*P. mexicana, P. latipinna, P. formosa*, Cyprinodontiformes, Poecilidae) reticulata). Bulletin of Fish Biology, 20, 71–85.

7. Buchanan, S. M., Kain, J. S., & De Bivort, B. L. (2015). Neuronal control of locomotor handedness in Drosophila. Proceedings of the National Academy of Sciences, 112(21), 6700–6705.

8. Chang, C., Moiron, M., Sánchez-Tójar, A., Niemelä, P. T., & Laskowski, K. L. (2024). What is the meta-analytic evidence for life-history trade-offs at the genetic level? Ecology Letters, 27(1). 10.1111/ele.14354

9. Clare, M. J., & Luckinbill, L. S. (1985). The effects of gene-environment interaction on the expression of longevity. Heredity, 55(1), 19–26. 10.1038/hdy.1985.67

10. Cressler, C. E., Bengtson, S., & Nelson, W. A. (2017). Unexpected nongenetic individual heterogeneity and trait covariance in *Daphnia* and its consequences for ecological and evolutionary dynamics. The American Naturalist, 190(1), E13–E27. 10.1086/691779

11. Dammhahn, M., Dingemanse, N. J., Niemelä, P. T., & Réale, D. (2018). Pace-of-life syndromes: a framework for the adaptive integration of behaviour, physiology and life history. Behavioral Ecology and Sociobiology, 72(3), 62. 10.1007/s00265-018-2473-y

12. Dönertaş, H. M., Fabian, D. K., Fuentealba, M., Partridge, L., & Thornton, J. M. (2021). Common genetic associations between age-related diseases. Nature Aging, 1(4), 400–412. 10.1038/s43587-021-00051-5

13. Duckworth, R. A. (2010). Evolution of personality: Developmental constraints on behavioral flexibility. The Auk, 127(4), 752–758. 10.1525/auk.2010.127.4.752

14. Ehlman, S. M., Scherer, U., Bierbach, D., Francisco, F. A., Laskowski, K. L., Krause, J., & Wolf, M. (2023). Leveraging big data to uncover the eco-evolutionary factors shaping behavioural development. Proceedings of the Royal Society B, 290, 20222115.

15. Freund, J., Brandmaier, A. M., Lewejohann, L., Kirste, I., Kritzler, M., Krüger, A., Sachser, N., Lindenberger, U., & Kempermann, G. (2013). Emergence of Individuality in Genetically Identical Mice. Science, 340(6133), 756–759. 10.1126/science.1235294

16. Herndon, L. A., Schmeissner, P. J., Dudaronek, J. M., Brown, P. A., Listner, K. M., Sakano, Y., Paupard, M. C., Hall, D. H., & Driscoll, M. (2002). Stochastic and genetic factors influence tissue-specific decline in ageing *C. elegans*. Nature, 419(6909), 808–814. 10.1038/nature01135

17. Hertel, A. G., Niemelä, P. T., Dingemanse, N. J., & Mueller, T. (2020). A guide for studying among-individual behavioral variation from movement data in the wild. Movement Ecology, 8(1), 30. 10.1186/s40462-020-00216-8

18. Johnson, J., Sheehy, K., & Laskowski, K. L. (2025). Playing dice with behavior: drivers of stochastic individuality. Trends in Ecology & Evolution. 10.1016/j.tree.2025.10.003

19. Kain, J. S., Stokes, C., & de Bivort, B. L. (2012). Phototactic personality in fruit flies and its suppression by serotonin and white. Proceedings of the National Academy of Sciences, 109(48), 19834–19839.

20. Kempermann, G. (2019). Environmental enrichment, new neurons and the neurobiology of individuality. Nature Reviews Neuroscience, 20(4), 235–245. 10.1038/s41583-019-0120-x

21. Kempermann, G., Lopes, J. B., Zocher, S., Schilling, S., Ehret, F., Garthe, A., Karasinsky, A., Brandmaier, A. M., Lindenberger, U., Winter, Y., & Overall, R. W. (2022). The individuality paradigm: Automated longitudinal activity tracking of large cohorts of genetically identical mice in an enriched environment. Neurobiology of Disease, 175, 105916. 10.1016/j.nbd.2022.105916

22. Kirkwood, T. B. L., Feder, M., Finch, C. E., Franceschi, C., Globerson, A., Klingenberg, C. P., LaMarco, K., Omholt, S., & Westendorp, R. G. J. (2005). What accounts for the wide variation in life span of genetically identical organisms reared in a constant environment? Mechanisms of Ageing and Development, 126(3), 439–443. 10.1016/j.mad.2004.09.008

23. Kolora, S. R. R., Owens, G. L., Vazquez, J. M., Stubbs, A., Chatla, K., Jainese, C., Seeto, K., McCrea, M., Sandel, M. W., Vianna, J. A., Maslenikov, K., Bachtrog, D., Orr, J. W., Love, M., & Sudmant, P. H. (2021). Origins and evolution of extreme life span in Pacific Ocean rockfishes. Science, 374(6569), 842–847. 10.1126/science.abg5332

24. Lamatsch, D. K., Schmid, M., & Schartl, M. (2005). A somatic mosaic of the gynogenetic Amazon molly. Journal of Fish Biology, 60(6), 1417–1422.

25. Lampert, K. P., & Schartl, M. (2008). The origin and evolution of a unisexual hybrid: *Poecilia formosa*. Philosophical Transactions of the Royal Society B: Biological Sciences, 363(1505), 2901–2909. 10.1098/rstb.2008.0040

26. Laskowski, K. L., Bierbach, D., Jolles, J. W., Doran, C., & Wolf, M. (2022). The emergence and development of behavioral individuality in clonal fish. Nature Communications, 13(1), 6419. 10.1038/s41467-022-34113-y

27. Linneweber, G. A., Andriatsilavo, M., Dutta, S. B., Bengochea, M., Hellbruegge, L., Liu, G., Ejsmont, R. K., Straw, A. D., Wernet, M., Hiesinger, P. R., & Hassan, B. A. (2020). A neurodevelopmental origin of behavioral individuality in the *Drosophila* visual system. Science, 367(6482), 1112–1119. 10.1126/science.aaw7182

28. Lüdecke, D. (2022). sjPlot: Data visualization for statistics in social science (R package version 2.8.12). https://strengejacke.github.io/sjPlot

29. Mönck, H. J., Jörg, A., von Falkenhausen, T., Tanke, J., Wild, B., Dormagen, D., Piotrowski, J., Winklmayr, C., Bierbach, D., & Landgraf, T. (2018). BioTracker: An open-source computer vision framework for visual animal tracking. *Computer Science*, arXiv:1803.07985.

30. Munch, S. B., & Salinas, S. (2009). Latitudinal variation in lifespan within species is explained by the metabolic theory of ecology. Proceedings of the National Academy of Sciences, 106(33), 13860–13864. 10.1073/pnas.0900300106

31. Niemelä, P. T., Vainikka, A., Forsman, J. T., Loukola, O. J., & Kortet, R. (2013). How does variation in the environment and individual cognition explain the existence of consistent behavioral differences? Ecology and Evolution, 3(2), 457–464. 10.1002/ece3.451

32. Poulain, M. (2013). The blue zones: areas of exceptional longevity around the sorld. Vienna Yearbood of Population Research, 87–108.

33. R Core Team. (2022). R: A language and environment for statistical computing. R Foundation for Statistical Computing.

34. Rea, S. L., Wu, D., Cypser, J. R., Vaupel, J. W., & Johnson, T. E. (2005). A stress-sensitive reporter predicts longevity in isogenic populations of Caenorhabditis elegans. Nature Genetics, 37(8), 894–898. 10.1038/ng1608

35. Réale, D., Garant, D., Humphries, M. M., Bergeron, P., Careau, V., & Montiglio, P.-O. (2010). Personality and the emergence of the pace-of-life syndrome concept at the population level. Philosophical Transactions of the Royal Society B: Biological Sciences, 365(1560), 4051–4063. 10.1098/rstb.2010.0208

36. Réale, D., Reader, S. M., Sol, D., McDougall, P. T., & Dingemanse, N. J. (2007). Integrating animal temperament within ecology and evolution. Biological Reviews, 82(2), 291–318. 10.1111/j.1469-185X.2007.00010.x

37. Royauté, R., Berdal, M. A., Garrison, C. R., & Dochtermann, N. A. (2018). Paceless life? A meta-analysis of the pace-of-life syndrome hypothesis. Behavioral Ecology and Sociobiology, 72(3), 64. 10.1007/s00265-018-2472-z

38. Scherer, U., Ehlman, S. M., Bierbach, D., Krause, J., & Wolf, M. (2023). Reproductive individuality of clonal fish raised in near-identical environments and its link to early-life behavioral individuality. Nature Communications, 14(1), 7652. 10.1038/s41467-023-43069-6

39. Schlupp, I. (2005). The evolutionary ecology of gynogenesis. Annual Review of Ecology, Evolution, and Systematics, 36(1), 399–417. 10.1146/annurev.ecolsys.36.102003.152629

40. Schlupp, I. (2010). Mate choice and the Amazon molly: how sexuality and unisexuality can coexist. Journal of Heredity, 101(Supplement 1), S55–S61. 10.1093/jhered/esq015

41. Schneider, C. A., Rasband, W. S., & Eliceiri, K. W. (2012). NIH Image to ImageJ: 25 years of image analysis. Nat Meth, 9(7), 671–675. 10.1038/nmeth.2089

42. Schuett, W., Dall, S. R. X., Kloesener, M. H., Baeumer, J., Beinlich, F., & Eggers, T. (2015). Life-history trade-offs mediate “personality” variation in two colour morphs of the pea aphid, *Acyrthosiphon pisum*. Journal of Animal Ecology, 84(1), 90–101. http://www.jstor.org/stable/24698459

43. Sih, A., Bell, A., & Johnson, J. C. (2004). Behavioral syndromes: an ecological and evolutionary overview. Trends in Ecology & Evolution, 19(7), 372–378. 10.1016/j.tree.2004.04.009

44. Stärk, L., Scherer, U., Ehlman, S. M., & Wolf, M. (2022). Tool for fish trajectory data visualization and analysis (1.0.0). https://github.com/UlrikeScherer/Fish-Tracking-Visualization

45. Steiner, U. K., Lenart, A., Ni, M., Chen, P., Song, X., Taddei, F., Vaupel, J. W., & Lindner, A. B. (2019). Two stochastic processes shape diverse senescence patterns in a single-cell organism. Evolution, 73(4), 847–857. 10.1111/evo.13708

46. Stöck, M., Lampert, K. P., Möller, D., Schlupp, I., & Schartl, M. (2010a). Monophyletic origin of multiple clonal lineages in an asexual fish (*Poecilia formosa*). Molecular Ecology, 19(23), 5204–5215. 10.1111/j.1365-294X.2010.04869.x

47. Stöck, M., Lampert, K. P., Möller, D., Schlupp, I., & Schartl, M. (2010b). Monophyletic origin of multiple clonal lineages in an asexual fish *(Poecilia formosa)*. Molecular Ecology, 19(23), 5204–5215. 10.1111/j.1365-294X.2010.04869.x

48. Stoffel, M. A., Nakagawa, S., & Schielzeth, H. (2021). partR2: partitioning R ^2^ in generalized linear mixed models. PeerJ, 9, e11414. 10.7717/peerj.11414

49. Vogt, G. (2015). Stochastic developmental variation, an epigenetic source of phenotypic diversity with far-reaching biological consequences. Journal of Biosciences, 40(1), 159–204. 10.1007/s12038-015-9506-8

50. von Bertalanffy, L. (1938). A quantitive theory of organic growth (inquires on growth laws. II). Human Biology, 10(2), 181–213. http://www.jstor.org/stable/41447359

51. Warren, W. C., García-Pérez, R., Xu, S., Lampert, K. P., Chalopin, D., Stöck, M., Loewe, L., Lu, Y., Kuderna, L., Minx, P., Montague, M. J., Tomlinson, C., Hillier, L. W., Murphy, D. N., Wang, J., Wang, Z., Garcia, C. M., Thomas, G. C. W., Volff, J.-N., … Schartl, M. (2018). Clonal polymorphism and high heterozygosity in the celibate genome of the Amazon molly. Nature Ecology & Evolution, 2(4), 669–679. 10.1038/s41559-018-0473-y

52. Werkhoven, Z., Bravin, A., Skutt-Kakaria, K., Reimers, P., Pallares, L. F., Ayroles, J., & de Bivort, B. L. (2021). The structure of behavioral variation within a genotype. ELife, 10. 10.7554/eLife.64988

53. Wickham, H. (2016). ggplot2: elegant graphics for data analysis. Springer-Verlag.

54. Wolf, M., & Weissing, F. J. (2012). Animal personalities: consequences for ecology and evolution. Trends in Ecology & Evolution, 27(8), 452–461.

55. Zhu, P., Liu, W., Zhang, X., Li, M., Liu, G., Yu, Y., Li, Z., Li, X., Du, J., Wang, X., Grueter, C. C., Li, M., & Zhou, X. (2023). Correlated evolution of social organization and lifespan in mammals. Nature Communications, 14(1), 372. 10.1038/s41467-023-35869-7

56. Zipple, M. N., Chang Kuo, D., Meng, X., Reichard, T. M., Guess, K., Vogt, C. C., Moeller, A. H., & Sheehan, M. J. (2025). Competitive social feedback amplifies the role of early life contingency in male mice. Science, 387(6729), 81–85. 10.1126/science.adq0579

