## Supplementary Information for "Activity during the first days of life predicts lifespan in a naturally clonal vertebrate"

#### Supplementary Note 1: Additional figures

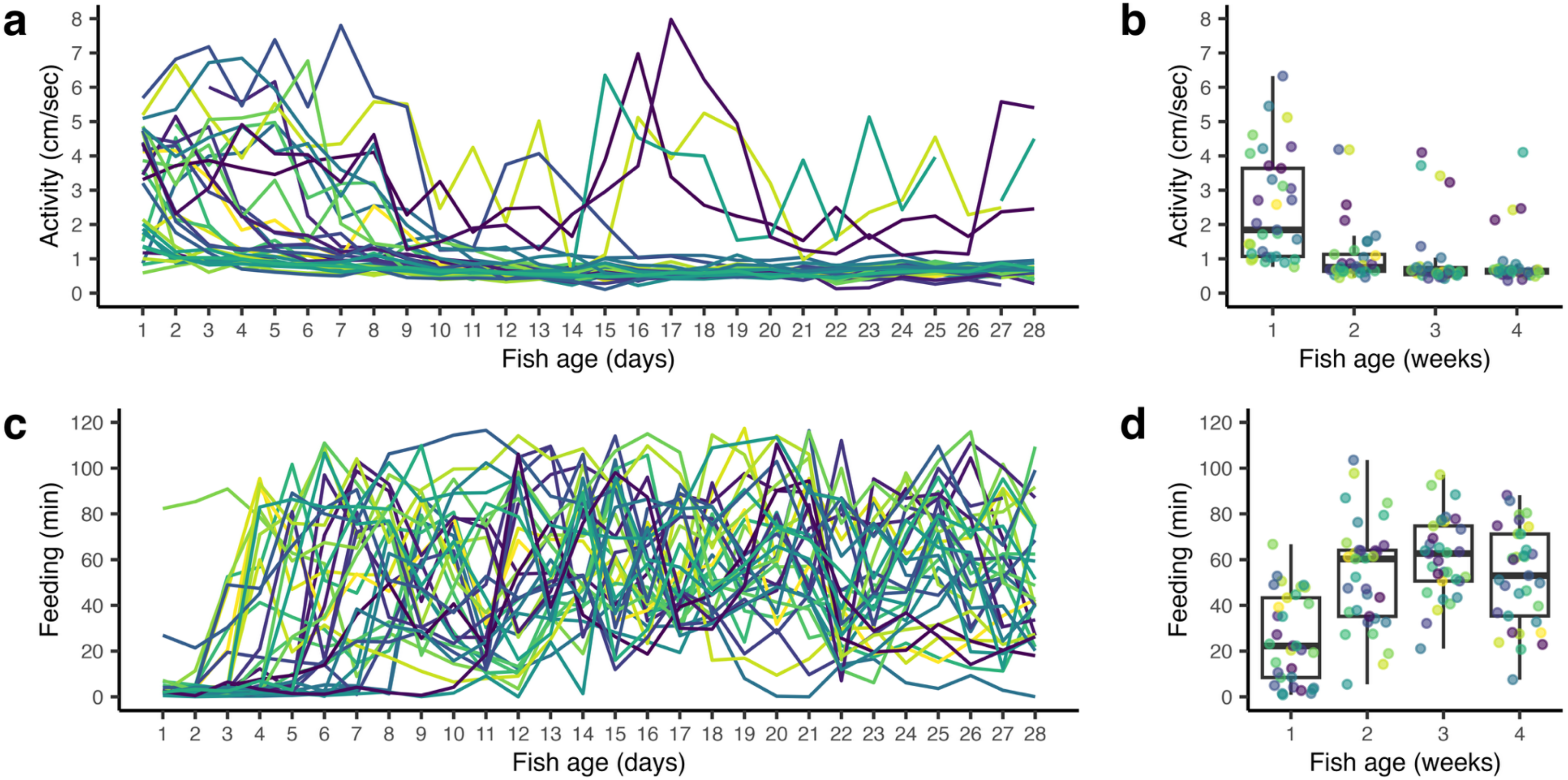

**Supplementary Figure 1. Behavioral variation expressed by genetically identical Amazon mollies, kept under near-identical conditions.** (a) Daily activity and (c) feeding behavior expressed by each individual (*N* = 33 individuals, respectively) over the first four weeks of their life. (b) Average activity and (d) feeding behavior expressed during each of the four observation weeks separately. (b, d) Boxplots with raw data (points, *N* = 33 data points per week, i.e., one data point per individual and week), medians (horizontal lines), 25% and 75% quantiles (boxes), and 1.5x interquartile ranges (whiskers). (a-d) Line and point color corresponds to fish ID, consistent across panels.

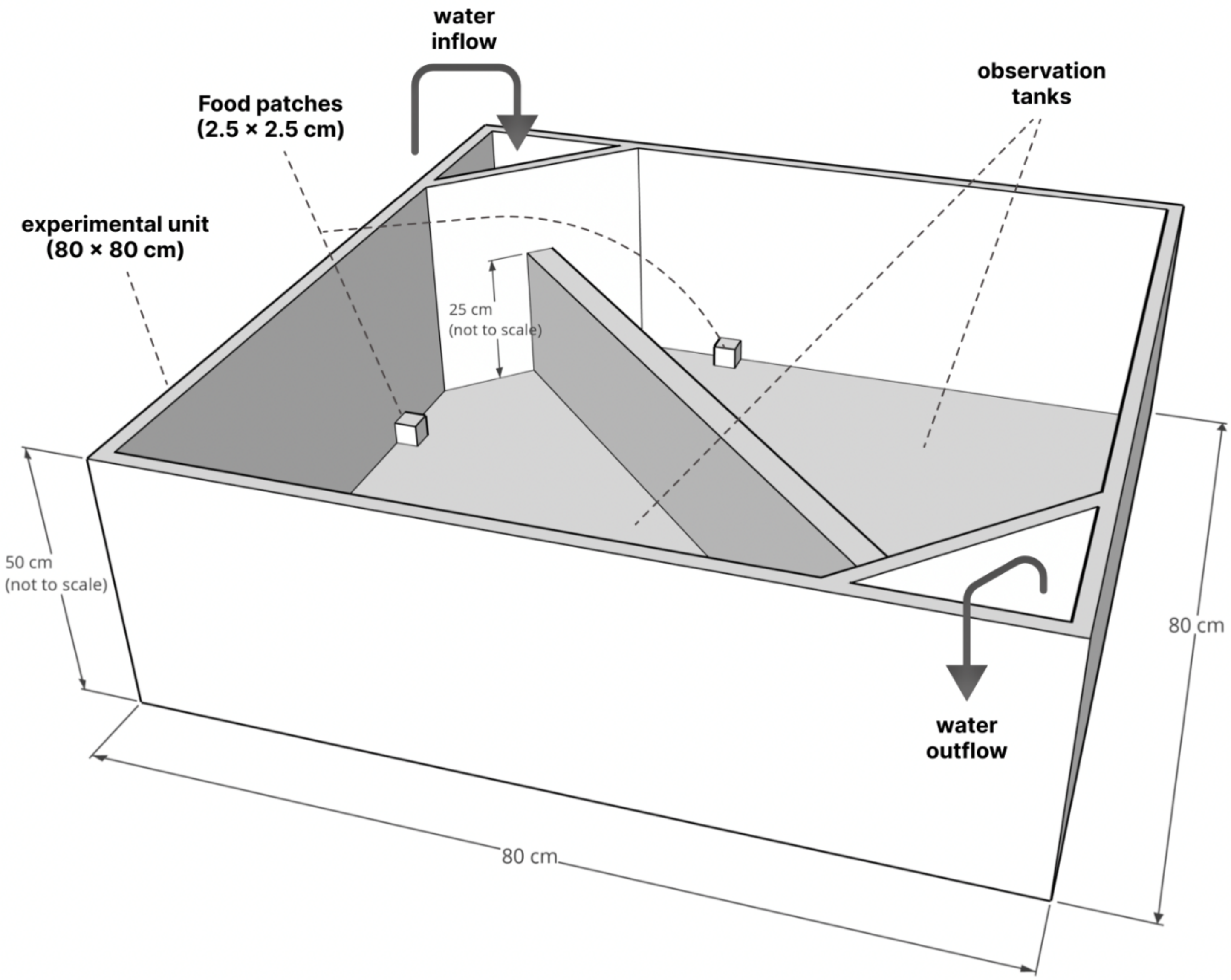

**Supplementary Figure 2**. "**Experimental set-up for behavioral observations (***Figure and legend adopted from Scherer et al. (2023):* "One experimental unit with 2 observation tanks. Water level in the tanks: 7 cm. Food patches were present during the feeding only. Observation tanks were illuminated individually from below with 4 LEDs per tank (each LED is 100cm, 12V, color temperature = 5500 K, light output = approx. 1570 lumen; tanks were manufactured from white polyethylene, which allowed light from underneath to get through). There was no visual contact between observation tanks, but tanks were connected via a flow-through water system (24 observation tanks split into 4 flow-through systems)."

### Supplementary Note 2: Model summaries main analyses

**Supplementary Table 1. Summary of linear mixed-effects models testing whether early-life behavior predicts lifespan.** Models assess the effects of activity and feeding behavior on lifespan when both behaviors are averaged across the first four weeks of life, and when analyses are restricted to each week separately to evaluate the temporal development of behavior–lifespan relationships.

| **Time interval** | **Response** | **Predictor** | **Full model** | | | | | **Final model** | | | | |
|---|---|---|---|---|---|---|---|---|---|---|---|---|
| | | | **Estimate** | **SE** | **$\chi^2$** | **p** | **df** | **Estimate** | **SE** | **$\chi^2$** | **p** | **df** |
| Week 1-4 | Lifespan (days) | (Intercept) | 595.260 | 13.941 | - | - | - | 636.824 | 14.390 | - | - | - |
| | | Activity (cm/sec, log-transformed) | -135.304 | -3.716 | 11.535 | 0.001 | 1 | -142.148 | 36.375 | 12.550 | <0.001 | 1 |
| | | Squared Activity (cm/sec, log-transformed) | 132.775 | 3.051 | 8.200 | 0.004 | 1 | 129.667 | 44.109 | 7.676 | 0.006 | 1 |
| | | Feeding (min) | 0.822 | 1.032 | 1.048 | 0.306 | 1 | - | - | - | - | - |
| | | **Random Effects** | | | | | | | | | | |
| | | $\sigma^2$ | 3970.70 | | | | | 4098.82 | | | | |
| | | $\tau_{00}$ | 0.00 Mother ID | | | | | 0.00 Mother ID | | | | |
| | | *N* | 3 Mother ID | | | | | 3 Mother ID | | | | |
| | | Observations | 33 | | | | | 33 | | | | |
| | | Marginal $R^2$ / Conditional $R^2$ | 0.347 / NA | | | | | 0.325 / NA | | | | |
| | | AIC | 353.533 | | | | | 353.981 | | | | |

**Supplementary Table 1.** Continued.

| Time interval | Response | Predictor | Full model | | | | | Final model | | | | |
|---|---|---|---|---|---|---|---|---|---|---|---|---|
| | | | **Estimate** | **SE** | **$\chi^2$** | **p** | **df** | **Estimate** | **SE** | **$\chi^2$** | **p** | **df** |
| Week 1 | Lifespan (days) | (Intercept) | 674.543 | 32.275 | - | - | - | 699.570 | 17.412 | - | - | - |
| | | Activity (cm/sec, log-transformed) | -67.690 | 60.780 | 1.218 | 0.230 | 1 | -64.740 | 18.331 | 10.580 | 0.001 | 1 |
| | | Squared Activity (cm/sec, log-transformed) | 9.538 | 36.779 | 0.067 | 0.796 | 1 | - | - | - | - | - |
| | | Feeding (min) | 0.731 | 0.705 | 1.058 | 0.304 | 1 | - | - | - | - | - |
| | | Random Effects | | | | | | | | | | |
| | | $\sigma^2$ | 4204.29 | | | | | 4363.52 | | | | |
| | | $\tau_{00}$ | 0.00 Mother ID | | | | | 0.00 Mother ID | | | | |
| | | *N* | 3 Mother ID | | | | | 3 Mother ID | | | | |
| | | Observations | 33 | | | | | 33 | | | | |
| | | Marginal $R^2$ / Conditional $R^2$ | 0.307 / NA | | | | | 0.280 / NA | | | | |
| | | AIC | 356.276 | | | | | 363.785 | | | | |

**Supplementary Table 1.** Continued.

| Time interval | Response | Predictor | Full model | | | | | Final model | | | | |
|---|---|---|---|---|---|---|---|---|---|---|---|---|
| | | | Estimate | SE | $\chi^2$ | p | df | Estimate | SE | $\chi^2$ | p | df |
| Week 2 | Lifespan (days) | (Intercept) | 648.420 | 33.600 | - | - | - | 649.157 | 12.598 | - | - | - |
| | | Activity (cm/sec, log-transformed) | -94.235 | 29.386 | 8.952 | 0.003 | 1 | -52.860 | 22.221 | 5.223 | 0.022 | 1 |
| | | Squared Activity (cm/sec, log-transformed) | 60.795 | 32.314 | 3.362 | 0.077 | 1 | - | - | - | - | - |
| | | Feeding (min) | -0.414 | 0.560 | 0.543 | 0.461 | 1 | - | - | - | - | - |
| | | **Random Effects** | | | | | | | | | | |
| | | $\sigma^2$ | 4567.50 | | | | | 5132.66 | | | | |
| | | $\tau_{00}$ | 0.00 Mother ID | | | | | 0.00 Mother ID | | | | |
| | | *N* | 3 Mother ID | | | | | 3 Mother ID | | | | |
| | | Observations | 33 | | | | | 33 | | | | |
| | | Marginal $R^2$ / Conditional $R^2$ | 0.246 / NA | | | | | 0.150 / NA | | | | |
| | | AIC | 359.321 | | | | | 368.595 | | | | |

**Supplementary Table 1.** Continued.

| Time interval | Response | Predictor | Full model | | | | | Final model | | | | |
|---|---|---|---|---|---|---|---|---|---|---|---|---|
| | | | **Estimate** | **SE** | **$\chi^2$** | **p** | **df** | **Estimate** | **SE** | **$\chi^2$** | **p** | **df** |
| Week 3 | Lifespan (days) | (Intercept) | 496.801 | 52.632 | - | - | - | 588.510 | 27.139 | - | - | - |
| | | Activity (cm/sec, log-transformed) | -73.008 | 28.698 | 5.911 | 0.015 | 1 | -65.485 | 30.116 | 4.419 | 0.036 | 1 |
| | | Squared Activity (cm/sec, log-transformed) | 117.526 | 37.035 | 8.789 | 0.003 | 1 | 103.906 | 38.533 | 6.571 | 0.010 | 1 |
| | | Feeding (min) | 1.367 | 0.685 | 3.758 | 0.053 | 1 | - | - | - | - | - |
| | | **Random Effects** | | | | | | | | | | |
| | | $\sigma^2$ | 4391.91 | | | | | 4921.67 | | | | |
| | | $\tau_{00}$ | 0.00 Mother ID | | | | | 0.00 Mother ID | | | | |
| | | *N* | 3 Mother ID | | | | | 3 Mother ID | | | | |
| | | Observations | 33 | | | | | 33 | | | | |
| | | Marginal $R^2$ / Conditional $R^2$ | 0.276 / NA | | | | | 0.186 / NA | | | | |
| | | AIC | 357.657 | | | | | 360.268 | | | | |

**Supplementary Table 1.** Continued.

| Time interval | Response | Predictor | Full model | | | | | Final model | | | | |
|---|---|---|---|---|---|---|---|---|---|---|---|---|
| | | | Estimate | SE | $\chi^2$ | p | df | Estimate | SE | $\chi^2$ | p | df |
| Week 4 | Lifespan (days) | (Intercept) | 622.251 | 37.445 | - | - | - | 653.394 | 13.498 | - | - | - |
| | | Activity (cm/sec, log-transformed) | 8.139 | 33.775 | 0.058 | 0.811 | 1 | - | - | - | - | - |
| | | Squared Activity (cm/sec, log-transformed) | 16.745 | 44.388 | 0.142 | 0.706 | 1 | - | - | - | - | - |
| | | Feeding (min) | 0.524 | 0.711 | 0.541 | 0.463 | 1 | - | - | - | - | - |
| | | **Random Effects** | | | | | | | | | | |
| | | $\sigma^2$ | 5826.20 | | | | | 6012.78 | | | | |
| | | $\tau_{00}$ | 0.00 Mother ID | | | | | 0.00 Mother ID | | | | |
| | | *N* | 3 Mother ID | | | | | 3 Mother ID | | | | |
| | | Observations | 33 | | | | | 33 | | | | |
| | | Marginal $R^2$ / Conditional $R^2$ | 0.032 / NA | | | | | 0.000 / NA | | | | |
| | | AIC | 365.718 | | | | | 379.746 | | | | |

**Supplementary Table 2. Summary of a linear mixed-effects model testing for potential tank effects.** The model tests whether the spatial position of long-term housing tanks influences individual lifespan, with tank position being defined by vertical level (top, middle, bottom) and centrality (central vs. peripheral). Neither tank level nor centrality had a significant effect on lifespan, indicating that spatial heterogeneity in the housing setup did not confound longevity outcomes.

| Response | Predictor | Full model | | | | | Final model | | | | |
|---|---|---|---|---|---|---|---|---|---|---|---|
| | | **Estimate** | **SE** | **$\chi^2$** | **p** | **df** | **Estimate** | **SE** | **$\chi^2$** | **p** | **df** |
| Lifespan (days) | (Intercept) | 629.949 | 23.333 | - | - | - | 653.394 | 13.498 | - | - | - |
| | Tank level [middle] | 22.727 | 31.839 | 0.654 | 0.721 | 1 | - | - | - | - | - |
| | Tank level [top] | 22.091 | 31.839 | | | | - | - | - | - | - |
| | Tank centrality [periphery] | 46.778 | 33.701 | 1.872 | 0.171 | 1 | - | - | - | - | - |
| | **Random Effects** | | | | | | | | | | |
| | $\sigma^2$ | 5575.61 | | | | | 6012.78 | | | | |
| | $\tau_{00}$ | 0.00 Mother ID | | | | | 0.00 Mother ID | | | | |
| | *N* | 3 Mother ID | | | | | 3 Mother ID | | | | |
| | Observations | 33 | | | | | 33 | | | | |
| | Marginal $R^2$ / Conditional $R^2$ | 0.075 / NA | | | | | 0.000 / NA | | | | |
| | AIC | 356.989 | | | | | 379.746 | | | | |

**Supplementary Table 3. Overview experimental phases.** In addition to the focal analyses of early-life behavior and lifespan, individuals were characterized for additional phenotypic traits, including reproductive output (Scherer et al., 2023) and additional behavioral assays, consistent with 3R guidelines to maximize data collected from individual animals.

| Experimental phase | | Start age (day) | End age (days) | Duration (days) |
|---|---|---|---|---|
| Phase 1 | Behavioral typing I | 1 | 28 | 28 |
| Phase 2 | Reproduction I | 29 | 280 | 241 |
| Phase 3 | Individual housing (including Behavioral typing II) | 281 | Mom 1 & 2 = 445, Mom 3 = 393 | Mom 1 & 2 = 163, Mom 3 = 111 |
| | Behavioral typing II | Mom 1 & 2 = 369, Mom 3 = 371 | Mom 1 & 2 = 379, Mom 3 = 381 | 10 |
| Phase 4 | Reproduction II | Mom 1 & 2 = 445, Mom 3 = 393 | Mom 1 & 2 = 599, Mom 3 = 547 | 154 |
| Phase 5 | Individual housing | Mom 1 & 2 = 600, Mom 3 = 548 | - | - |

#### Supplementary Note 3: Daily analysis

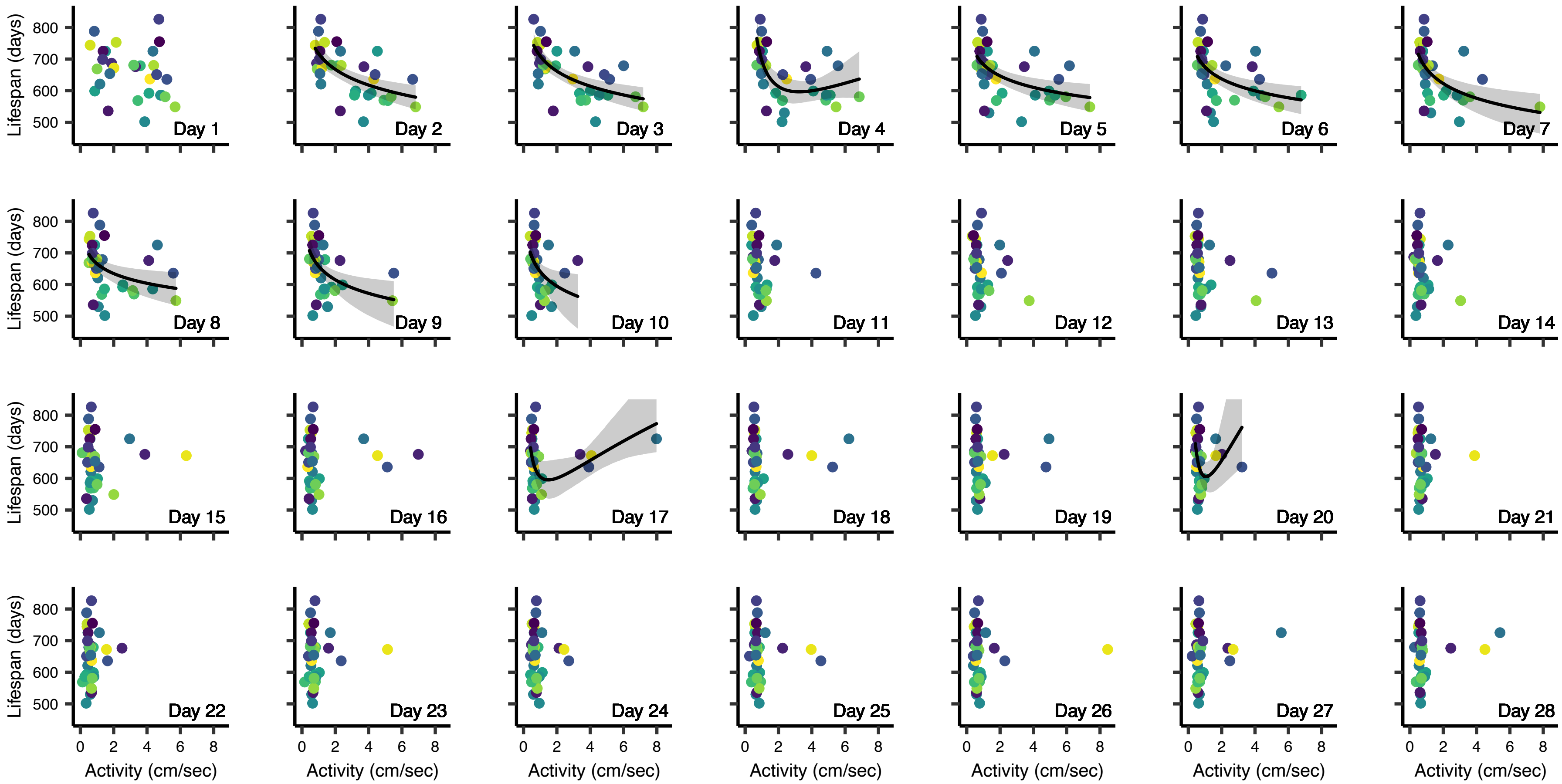

**Supplementary Figure 3. Daily resolution of the early-life activity–lifespan relationship.** Activity expressed on individual days during the first 28 days of life is plotted against lifespan. Each panel shows one day (day 1–28), with regression lines (black) and 95% confidence intervals (gray) shown only where the activity–lifespan relationship was statistically significant. Consistent with the weekly analyses, negative associations are most pronounced during the earliest days of life (day 2-10, except day 4), whereas U-shaped relationships emerge mostly later – when variation in activity is reduced (day 4, 17, and 20). Points represent raw data; activity values were log-transformed for statistical analyses and back-transformed for visualization. Point color indicates fish identity and is consistent across panels.

**Supplementary Table 4. Summary of linear mixed-effects models testing whether early-life behavior predicts lifespan at daily resolution.** Models were fitted separately for each of the first 28 days of life, i.e., behavioral predictors (activity and feeding) were restricted to the respective day only (i.e., one model per day; 28 models total). This analysis assesses the robustness and fine-scale temporal dynamics of the early-life behavior–lifespan relationship observed at broader time scales.

| **Time interval** | **Response** | **Predictor** | **Full model** | | | | | **Final model** | | | | |
|---|---|---|---|---|---|---|---|---|---|---|---|---|
| | | | **Estimate** | **SE** | **$\chi^2$** | **p** | **df** | **Estimate** | **SE** | **$\chi^2$** | **p** | **df** |
| Day 1 | Lifespan (days) | (Intercept) | 677.786 | 30.110 | - | - | - | 658.862 | 14.392 | - | - | - |
| | | Activity (cm/sec, log-transformed) | -9.903 | 77.986 | 0.016 | 0.899 | 1 | - | - | - | - | - |
| | | Squared Activity (cm/sec, log-transformed) | -11.273 | 46.310 | 0.059 | 0.808 | 1 | - | - | - | - | - |
| | | Feeding (min) | 0.871 | 1.177 | 0.542 | 0.462 | 1 | - | - | - | - | - |
| | | **Random Effects** | | | | | | | | | | |
| | | $\sigma^2$ | 5392.00 | | | | | 6007.15 | | | | |
| | | $\tau_{00}$ | 0.00 Mother ID | | | | | 0.00 Mother ID | | | | |
| | | *N* | 3 Mother ID | | | | | 3 Mother ID | | | | |
| | | Observations | 29 | | | | | 29 | | | | |
| | | Marginal $R^2$ / Conditional $R^2$ | 0.106 / NA | | | | | 0.000 / NA | | | | |
| | | AIC | 316.972 | | | | | 333.430 | | | | |

**Supplementary Table 4.** Continued.

| Time interval | Response | Predictor | Full model | | | | | Final model | | | | |
|---|---|---|---|---|---|---|---|---|---|---|---|---|
| | | | Estimate | SE | $\chi^2$ | p | df | Estimate | SE | $\chi^2$ | p | df |
| Day 2 | Lifespan (days) | (Intercept) | 706.349 | 21.699 | - | - | - | 715.838 | 17.251 | - | - | - |
| | | Activity (cm/sec, log-transformed) | -47.322 | 57.746 | 0.664 | 0.415 | 1 | -70.745 | 16.053 | 15.097 | <0.001 | 1 |
| | | Squared Activity (cm/sec, log-transformed) | -11.866 | 32.893 | 0.131 | 0.719 | 1 | - | - | - | - | - |
| | | Feeding (min) | 0.584 | 0.793 | 0.538 | 0.463 | 1 | - | - | - | - | - |
| | | **Random Effects** | | | | | | | | | | |
| | | $\sigma^2$ | 3549.92 | | | | | 3614.21 | | | | |
| | | $\tau_{00}$ | 0.00 Mother ID | | | | | 0.00 Mother ID | | | | |
| | | *N* | 3 Mother ID | | | | | 3 Mother ID | | | | |
| | | Observations | 31 | | | | | 31 | | | | |
| | | Marginal $R^2$ / Conditional $R^2$ | 0.404 / NA | | | | | 0.393 / NA | | | | |
| | | AIC | 329.074 | | | | | 335.894 | | | | |

**Supplementary Table 4.** Continued.

| Time interval | Response | Predictor | Full model | | | | | Final model | | | | |
|---|---|---|---|---|---|---|---|---|---|---|---|---|
| | | | **Estimate** | **SE** | **$\chi^2$** | **p** | **df** | **Estimate** | **SE** | **$\chi^2$** | **p** | **df** |
| Day 3 | Lifespan (days) | (Intercept) | 692.357 | 17.290 | - | - | - | 709.095 | 14.285 | - | - | - |
| | | Activity (cm/sec, log-transformed) | -96.469 | 38.255 | 5.800 | 0.016 | 1 | -68.309 | 13.498 | 18.816 | <0.001 | 1 |
| | | Squared Activity (cm/sec, log-transformed) | 24.117 | 23.259 | 1.057 | 0.304 | 1 | - | - | - | - | - |
| | | Feeding (min) | 0.797 | 0.477 | 2.675 | 0.102 | 1 | - | - | - | - | - |
| | | **Random Effects** | | | | | | | | | | |
| | | $\sigma^2$ | 2842.05 | | | | | 3171.58 | | | | |
| | | $\tau_{00}$ | 0.00 Mother ID | | | | | 0.00 Mother ID | | | | |
| | | *N* | 3 Mother ID | | | | | 3 Mother ID | | | | |
| | | Observations | 32 | | | | | 32 | | | | |
| | | Marginal $R^2$ / Conditional $R^2$ | 0.510 / NA | | | | | 0.452 / NA | | | | |
| | | AIC | 335.280 | | | | | 343.255 | | | | |

**Supplementary Table 4.** Continued.

| Time interval | Response | Predictor | Full model | | | | | Final model | | | | |
|---|---|---|---|---|---|---|---|---|---|---|---|---|
| | | | Estimate | SE | $\chi^2$ | p | df | Estimate | SE | $\chi^2$ | p | df |
| Day 4 | Lifespan (days) | (Intercept) | 684.883 | 21.034 | - | - | - | 694.277 | 14.023 | - | - | - |
| | | Activity (cm/sec, log-transformed) | -155.345 | 48.331 | 8.988 | 0.003 | 1 | -163.074 | 46.813 | 10.334 | 0.001 | 1 |
| | | Squared Activity (cm/sec, log-transformed) | 66.204 | 29.230 | 4.768 | 0.029 | 1 | 68.046 | 29.223 | 5.021 | 0.025 | 1 |
| | | Feeding (min) | 0.214 | 0.358 | 0.354 | 0.552 | 1 | - | - | - | - | - |
| | | **Random Effects** | | | | | | | | | | |
| | | $\sigma^2$ | 3579.26 | | | | | 3617.88 | | | | |
| | | $\tau_{00}$ | 0.00 Mother ID | | | | | 0.00 Mother ID | | | | |
| | | *N* | 3 Mother ID | | | | | 3 Mother ID | | | | |
| | | Observations | 33 | | | | | 33 | | | | |
| | | Marginal $R^2$ / Conditional $R^2$ | 0.412 / NA | | | | | 0.406 / NA | | | | |
| | | AIC | 353.344 | | | | | 351.563 | | | | |

**Supplementary Table 4.** Continued.

| Time interval | Response | Predictor | Full model | | | | | Final model | | | | |
|---|---|---|---|---|---|---|---|---|---|---|---|---|
| | | | **Estimate** | **SE** | **$\chi^2$** | **p** | **df** | **Estimate** | **SE** | **$\chi^2$** | **p** | **df** |
| Day 5 | Lifespan (days) | (Intercept) | 673.034 | 23.117 | - | - | - | 683.783 | 14.397 | - | - | - |
| | | Activity (cm/sec, log-transformed) | -97.729 | 46.559 | 4.136 | 0.042 | 1 | -52.556 | 14.949 | 10.498 | 0.001 | 1 |
| | | Squared Activity (cm/sec, log-transformed) | 32.596 | 27.265 | 1.399 | 0.237 | 1 | - | - | - | - | - |
| | | Feeding (min) | 0.195 | 0.399 | 0.238 | 0.626 | 1 | - | - | - | - | - |
| | | **Random Effects** | | | | | | | | | | |
| | | $\sigma^2$ | 4119.57 | | | | | 4374.44 | | | | |
| | | $\tau_{00}$ | 0.00 Mother ID | | | | | 0.00 Mother ID | | | | |
| | | *N* | 3 Mother ID | | | | | 3 Mother ID | | | | |
| | | Observations | 33 | | | | | 33 | | | | |
| | | Marginal $R^2$ / Conditional $R^2$ | 0.322 / NA | | | | | 0.279 / NA | | | | |
| | | AIC | 357.824 | | | | | 364.273 | | | | |

**Supplementary Table 4.** Continued.

| Time interval | Response | Predictor | Full model | | | | | Final model | | | | |
|---|---|---|---|---|---|---|---|---|---|---|---|---|
| | | | Estimate | SE | $\chi^2$ | p | df | Estimate | SE | $\chi^2$ | p | df |
| Day 6 | Lifespan (days) | (Intercept) | 666.914 | 22.842 | - | - | - | 675.187 | 13.442 | - | - | - |
| | | Activity (cm/sec, log-transformed) | -114.536 | 39.240 | 7.579 | 0.007 | 1 | -53.566 | 16.241 | 9.402 | 0.002 | 1 |
| | | Squared Activity (cm/sec, log-transformed) | 48.239 | 26.944 | 3.059 | 0.085 | 1 | - | - | - | - | - |
| | | Feeding (min) | 0.001 | 0.364 | 0.000 | 0.998 | 1 | - | - | - | - | - |
| | | **Random Effects** | | | | | | | | | | |
| | | $\sigma^2$ | 4116.77 | | | | | 4522.09 | | | | |
| | | $\tau_{00}$ | 0.00 Mother ID | | | | | 0.00 Mother ID | | | | |
| | | *N* | 3 Mother ID | | | | | 3 Mother ID | | | | |
| | | Observations | 33 | | | | | 33 | | | | |
| | | Marginal $R^2$ / Conditional $R^2$ | 0.322 / NA | | | | | 0.254 / NA | | | | |
| | | AIC | 357.837 | | | | | 365.169 | | | | |

**Supplementary Table 4.** Continued.

| Time interval | Response | Predictor | Full model | | | | | Final model | | | | |
|---|---|---|---|---|---|---|---|---|---|---|---|---|
| | | | Estimate | SE | $\chi^2$ | p | df | Estimate | SE | $\chi^2$ | p | df |
| Day 7 | Lifespan (days) | (Intercept) | 677.727 | 22.808 | - | - | - | 668.963 | 12.346 | - | - | - |
| | | Activity (cm/sec, log-transformed) | -101.236 | 32.079 | 8.671 | 0.003 | 1 | -65.353 | 17.331 | 11.765 | 0.001 | 1 |
| | | Squared Activity (cm/sec, log-transformed) | 27.789 | 24.026 | 1.311 | 0.252 | 1 | - | - | - | - | - |
| | | Feeding (min) | -0.284 | 0.347 | 0.662 | 0.416 | 1 | - | - | - | - | - |
| | | **Random Effects** | | | | | | | | | | |
| | | $\sigma^2$ | 4030.36 | | | | | 4281.69 | | | | |
| | | $\tau_{00}$ | 0.00 Mother ID | | | | | 0.00 Mother ID | | | | |
| | | *N* | 3 Mother ID | | | | | 3 Mother ID | | | | |
| | | Observations | 32 | | | | | 32 | | | | |
| | | Marginal $R^2$ / Conditional $R^2$ | 0.356 / NA | | | | | 0.314 / NA | | | | |
| | | AIC | 346.130 | | | | | 352.058 | | | | |

**Supplementary Table 4.** Continued.

| Time interval | Response | Predictor | Full model | | | | | Final model | | | | |
|---|---|---|---|---|---|---|---|---|---|---|---|---|
| | | | Estimate | SE | $\chi^2$ | p | df | Estimate | SE | $\chi^2$ | p | df |
| Day 8 | Lifespan (days) | (Intercept) | 641.062 | 29.107 | - | - | - | 665.878 | 13.360 | - | - | - |
| | | Activity (cm/sec, log-transformed) | -85.604 | 34.629 | 5.607 | 0.018 | 1 | -43.320 | 17.389 | 5.687 | 0.018 | 1 |
| | | Squared Activity (cm/sec, log-transformed) | 42.574 | 27.183 | 2.366 | 0.124 | 1 | - | - | - | - | - |
| | | Feeding (min) | 0.225 | 0.422 | 0.283 | 0.595 | 1 | - | - | - | - | - |
| | | **Random Effects** | | | | | | | | | | |
| | | $\sigma^2$ | 4685.10 | | | | | 5061.01 | | | | |
| | | $\tau_{00}$ | 0.00 Mother ID | | | | | 0.00 Mother ID | | | | |
| | | *N* | 3 Mother ID | | | | | 3 Mother ID | | | | |
| | | Observations | 33 | | | | | 33 | | | | |
| | | Marginal $R^2$ / Conditional $R^2$ | 0.226 / NA | | | | | 0.162 / NA | | | | |
| | | AIC | 361.500 | | | | | 368.635 | | | | |

**Supplementary Table 4.** Continued.

| *Time interval* | *Response* | *Predictor* | **Full model** | | | | | **Final model** | | | | |
|---|---|---|---|---|---|---|---|---|---|---|---|---|
| | | | ***Estimate*** | ***SE*** | **$\chi^2$** | ***p*** | ***df*** | ***Estimate*** | ***SE*** | **$\chi^2$** | ***p*** | ***df*** |
| Day 9 | Lifespan (days) | (Intercept) | 656.437 | 26.830 | - | - | - | 658.519 | 12.293 | - | - | - |
| | | Activity (cm/sec, log-transformed) | -87.463 | 32.227 | 6.649 | 0.010 | 1 | -58.985 | 21.246 | 6.927 | 0.009 | 1 |
| | | Squared Activity (cm/sec, log-transformed) | 30.136 | 26.704 | 1.251 | 0.264 | 1 | - | - | - | - | - |
| | | Feeding (min) | -0.105 | 0.422 | 0.061 | 0.804 | 1 | - | - | - | - | - |
| | | **Random Effects** | | | | | | | | | | |
| | | $\sigma^2$ | 4683.76 | | | | | 4874.31 | | | | |
| | | $\tau_{00}$ | 0.00 Mother ID | | | | | 0.00 Mother ID | | | | |
| | | *N* | 3 Mother ID | | | | | 3 Mother ID | | | | |
| | | Observations | 33 | | | | | 33 | | | | |
| | | Marginal $R^2$ / Conditional $R^2$ | 0.226 / NA | | | | | 0.194 / NA | | | | |
| | | AIC | 361.084 | | | | | 367.032 | | | | |

**Supplementary Table 4.** Continued.

| Time interval | Response | Predictor | Full model | | | | | Final model | | | | |
|---|---|---|---|---|---|---|---|---|---|---|---|---|
| | | | Estimate | SE | $\chi^2$ | p | df | Estimate | SE | $\chi^2$ | p | df |
| Day 10 | Lifespan (days) | (Intercept) | 646.051 | 25.144 | - | - | - | 643.065 | 13.164 | - | - | - |
| | | Activity (cm/sec, log-transformed) | -75.282 | 24.649 | 8.215 | 0.004 | 1 | -61.302 | 25.434 | 5.351 | 0.020 | 1 |
| | | Squared Activity (cm/sec, log-transformed) | 75.989 | 39.866 | 3.447 | 0.063 | 1 | - | - | - | - | - |
| | | Feeding (min) | -0.495 | 0.394 | 1.547 | 0.214 | 1 | - | - | - | - | - |
| | | **Random Effects** | | | | | | | | | | |
| | | $\sigma^2$ | 4449.57 | | | | | 5112.73 | | | | |
| | | $\tau_{00}$ | 0.00 Mother ID | | | | | 0.00 Mother ID | | | | |
| | | *N* | 3 Mother ID | | | | | 3 Mother ID | | | | |
| | | Observations | 33 | | | | | 33 | | | | |
| | | Marginal $R^2$ / Conditional $R^2$ | 0.266 / NA | | | | | 0.154 / NA | | | | |
| | | AIC | 358.526 | | | | | 368.200 | | | | |

**Supplementary Table 4.** Continued.

| *Time interval* | *Response* | *Predictor* | **Full model** | | | | | **Final model** | | | | |
|---|---|---|---|---|---|---|---|---|---|---|---|---|
| | | | *Estimate* | *SE* | $\chi^2$ | *p* | *df* | *Estimate* | *SE* | $\chi^2$ | *p* | *df* |
| Day 11 | Lifespan (days) | (Intercept) | 626.658 | 25.023 | - | - | - | 641.231 | 14.530 | - | - | - |
| | | Activity (cm/sec, log-transformed) | -55.610 | 24.969 | 4.621 | 0.032 | 1 | -46.152 | 25.543 | 3.113 | 0.078 | 1 |
| | | Squared Activity (cm/sec, log-transformed) | 58.795 | 31.209 | 3.371 | 0.067 | 1 | - | - | - | - | - |
| | | Feeding (min) | -0.160 | 0.404 | 0.156 | 0.693 | 1 | - | - | - | - | - |
| | | **Random Effects** | | | | | | | | | | |
| | | $\sigma^2$ | 4871.62 | | | | | 5471.51 | | | | |
| | | $\tau_{00}$ | 0.00 Mother ID | | | | | 0.00 Mother ID | | | | |
| | | *N* | 3 Mother ID | | | | | 3 Mother ID | | | | |
| | | Observations | 33 | | | | | 33 | | | | |
| | | Marginal $R^2$ / Conditional $R^2$ | 0.195 / NA | | | | | 0.093 / NA | | | | |
| | | AIC | 361.854 | | | | | 370.362 | | | | |

**Supplementary Table 4.** Continued.

| *Time interval* | *Response* | *Predictor* | **Full model** | | | | | **Final model** | | | | |
|---|---|---|---|---|---|---|---|---|---|---|---|---|
| | | | ***Estimate*** | ***SE*** | **$\chi^2$** | ***p*** | ***df*** | ***Estimate*** | ***SE*** | **$\chi^2$** | ***p*** | ***df*** |
| Day 12 | Lifespan (days) | (Intercept) | 633.734 | 27.522 | - | - | - | 653.394 | 13.498 | - | - | - |
| | | Activity (cm/sec, log-transformed) | -45.409 | 23.808 | 3.451 | 0.063 | 1 | - | - | - | - | - |
| | | Squared Activity (cm/sec, log-transformed) | 37.436 | 33.741 | 1.209 | 0.272 | 1 | - | - | - | - | - |
| | | Feeding (min) | -0.117 | 0.359 | 0.105 | 0.746 | 1 | - | - | - | - | - |
| | | **Random Effects** | | | | | | | | | | |
| | | $\sigma^2$ | 5299.27 | | | | | 6012.78 | | | | |
| | | $\tau_{00}$ | 0.00 Mother ID | | | | | 0.00 Mother ID | | | | |
| | | *N* | 3 Mother ID | | | | | 3 Mother ID | | | | |
| | | Observations | 33 | | | | | 33 | | | | |
| | | Marginal $R^2$ / Conditional $R^2$ | 0.122 / NA | | | | | 0.000 / NA | | | | |
| | | AIC | 364.644 | | | | | 379.746 | | | | |

**Supplementary Table 4.** Continued.

| Time interval | Response | Predictor | Full model | | | | | Final model | | | | |
|---|---|---|---|---|---|---|---|---|---|---|---|---|
| | | | **Estimate** | **SE** | **$\chi^2$** | **p** | **df** | **Estimate** | **SE** | **$\chi^2$** | **p** | **df** |
| Day 13 | Lifespan (days) | (Intercept) | 623.964 | 35.891 | - | - | - | 653.394 | 13.498 | - | - | - |
| | | Activity (cm/sec, log-transformed) | -56.885 | 27.244 | -2.088 | 0.046 | 1 | - | - | - | - | - |
| | | Squared Activity (cm/sec, log-transformed) | 36.622 | 30.197 | 1.213 | 0.236 | 1 | - | - | - | - | - |
| | | Feeding (min) | -0.074 | 0.517 | -0.143 | 0.887 | 1 | - | - | - | - | - |
| | | **Random Effects** | | | | | | | | | | |
| | | $\sigma^2$ | 5307.47 | | | | | 6012.78 | | | | |
| | | $\tau_{00}$ | 0.00 Mother ID | | | | | 0.00 Mother ID | | | | |
| | | *N* | 3 Mother ID | | | | | 3 Mother ID | | | | |
| | | Observations | 33 | | | | | 33 | | | | |
| | | Marginal $R^2$ / Conditional $R^2$ | 0.121 / NA | | | | | 0.000 / NA | | | | |
| | | AIC | 364.373 | | | | | 379.746 | | | | |

**Supplementary Table 4.** Continued.

| Time interval | Response | Predictor | Full model | | | | | Final model | | | | |
|---|---|---|---|---|---|---|---|---|---|---|---|---|
| | | | Estimate | SE | $\chi^2$ | p | df | Estimate | SE | $\chi^2$ | p | df |
| Day 14 | Lifespan (days) | (Intercept) | 628.638 | 38.493 | - | - | - | 653.394 | 13.498 | - | - | - |
| | | Activity (cm/sec, log-transformed) | -27.883 | 27.522 | 1.011 | 0.315 | 1 | - | - | - | - | - |
| | | Squared Activity (cm/sec, log-transformed) | 6.121 | 32.094 | 0.036 | 0.849 | 1 | - | - | - | - | - |
| | | Feeding (min) | 0.126 | 0.492 | 0.066 | 0.798 | 1 | - | - | - | - | - |
| | | **Random Effects** | | | | | | | | | | |
| | | $\sigma^2$ | 5766.03 | | | | | 6012.78 | | | | |
| | | $\tau_{00}$ | 0.00 Mother ID | | | | | 0.00 Mother ID | | | | |
| | | *N* | 3 Mother ID | | | | | 3 Mother ID | | | | |
| | | Observations | 33 | | | | | 33 | | | | |
| | | Marginal $R^2$ / Conditional $R^2$ | 0.042 / NA | | | | | 0.000 / NA | | | | |
| | | AIC | 366.686 | | | | | 379.746 | | | | |

**Supplementary Table 4.** Continued.

| Time interval | Response | Predictor | Full model | | | | | Final model | | | | |
|---|---|---|---|---|---|---|---|---|---|---|---|---|
| | | | Estimate | SE | $\chi^2$ | p | df | Estimate | SE | $\chi^2$ | p | df |
| Day 15 | Lifespan (days) | (Intercept) | 635.689 | 33.304 | - | - | - | 653.394 | 13.498 | - | - | - |
| | | Activity (cm/sec, log-transformed) | -3.459 | 18.411 | 0.035 | 0.851 | 1 | - | - | - | - | - |
| | | Squared Activity (cm/sec, log-transformed) | 9.683 | 12.940 | 0.556 | 0.456 | 1 | - | - | - | - | - |
| | | Feeding (min) | 0.171 | 0.456 | 0.139 | 0.709 | 1 | - | - | - | - | - |
| | | **Random Effects** | | | | | | | | | | |
| | | $\sigma^2$ | 5886.02 | | | | | 6012.78 | | | | |
| | | $\tau_{00}$ | 0.00 Mother ID | | | | | 0.00 Mother ID | | | | |
| | | *N* | 3 Mother ID | | | | | 3 Mother ID | | | | |
| | | Observations | 33 | | | | | 33 | | | | |
| | | Marginal $R^2$ / Conditional $R^2$ | 0.022 / NA | | | | | 0.000 / NA | | | | |
| | | AIC | 369.947 | | | | | 379.746 | | | | |

**Supplementary Table 4.** Continued.

| Time interval | Response | Predictor | Full model | | | | | Final model | | | | |
|---|---|---|---|---|---|---|---|---|---|---|---|---|
| | | | Estimate | SE | $\chi^2$ | p | df | Estimate | SE | $\chi^2$ | p | df |
| Day 16 | Lifespan (days) | (Intercept) | 655.400 | 43.382 | - | - | - | 653.394 | 13.498 | - | - | - |
| | | Activity (cm/sec, log-transformed) | -10.161 | 21.311 | 0.227 | 0.634 | 1 | - | - | - | - | - |
| | | Squared Activity (cm/sec, log-transformed) | 16.160 | 20.582 | 0.611 | 0.435 | 1 | - | - | - | - | - |
| | | Feeding (min) | -0.308 | 0.534 | 0.331 | 0.565 | 1 | - | - | - | - | - |
| | | **Random Effects** | | | | | | | | | | |
| | | $\sigma^2$ | 5768.85 | | | | | 6012.78 | | | | |
| | | $\tau_{00}$ | 0.00 Mother ID | | | | | 0.00 Mother ID | | | | |
| | | *N* | 3 Mother ID | | | | | 3 Mother ID | | | | |
| | | Observations | 33 | | | | | 33 | | | | |
| | | Marginal $R^2$ / Conditional $R^2$ | 0.042 / NA | | | | | 0.000 / NA | | | | |
| | | AIC | 368.341 | | | | | 379.746 | | | | |

**Supplementary Table 4.** Continued.

| Time interval | Response | Predictor | Full model | | | | | Final model | | | | |
|---|---|---|---|---|---|---|---|---|---|---|---|---|
| | | | Estimate | SE | $\chi^2$ | p | df | Estimate | SE | $\chi^2$ | p | df |
| Day 17 | Lifespan (days) | (Intercept) | 572.632 | 42.780 | - | - | - | 608.718 | 25.038 | - | - | - |
| | | Activity (cm/sec, log-transformed) | -54.868 | 30.069 | 3.172 | 0.075 | 1 | -48.982 | 29.995 | 2.564 | 0.109 | 1 |
| | | Squared Activity (cm/sec, log-transformed) | 63.554 | 26.703 | 5.227 | 0.022 | 1 | 56.329 | 26.181 | 4.332 | 0.037 | 1 |
| | | Feeding (min) | 0.541 | 0.524 | 1.048 | 0.306 | 1 | - | - | - | - | - |
| | | **Random Effects** | | | | | | | | | | |
| | | $\sigma^2$ | 5104.97 | | | | | 5269.71 | | | | |
| | | $\tau_{00}$ | 0.00 Mother ID | | | | | 0.00 Mother ID | | | | |
| | | *N* | 3 Mother ID | | | | | 3 Mother ID | | | | |
| | | Observations | 33 | | | | | 33 | | | | |
| | | Marginal $R^2$ / Conditional $R^2$ | 0.155 / NA | | | | | 0.127 / NA | | | | |
| | | AIC | 363.812 | | | | | 363.424 | | | | |

**Supplementary Table 4.** Continued.

| Time interval | Response | Predictor | Full model | | | | | Final model | | | | |
|---|---|---|---|---|---|---|---|---|---|---|---|---|
| | | | Estimate | SE | $\chi^2$ | p | df | Estimate | SE | $\chi^2$ | p | df |
| Day 18 | Lifespan (days) | (Intercept) | 577.369 | 46.365 | - | - | - | 653.394 | 13.498 | - | - | - |
| | | Activity (cm/sec, log-transformed) | -53.928 | 35.428 | 2.239 | 0.135 | 1 | - | - | - | - | - |
| | | Squared Activity (cm/sec, log-transformed) | 67.064 | 33.230 | 3.841 | 0.050 | 1 | - | - | - | - | - |
| | | Feeding (min) | 0.451 | 0.564 | 0.634 | 0.426 | 1 | - | - | - | - | - |
| | | **Random Effects** | | | | | | | | | | |
| | | $\sigma^2$ | 5319.12 | | | | | 6012.78 | | | | |
| | | $\tau_{00}$ | 0.00 Mother ID | | | | | 0.00 Mother ID | | | | |
| | | *N* | 3 Mother ID | | | | | 3 Mother ID | | | | |
| | | Observations | 33 | | | | | 33 | | | | |
| | | Marginal $R^2$ / Conditional $R^2$ | 0.119 / NA | | | | | 0.000 / NA | | | | |
| | | AIC | 364.365 | | | | | 379.746 | | | | |

**Supplementary Table 4.** Continued.

| Time interval | Response | Predictor | Full model | | | | | Final model | | | | |
|---|---|---|---|---|---|---|---|---|---|---|---|---|
| | | | **Estimate** | **SE** | **$\chi^2$** | **p** | **df** | **Estimate** | **SE** | **$\chi^2$** | **p** | **df** |
| Day 19 | Lifespan (days) | (Intercept) | 580.484 | 45.437 | - | - | - | 652.750 | 14.140 | - | - | - |
| | | Activity (cm/sec, log-transformed) | -39.235 | 35.290 | 1.205 | 0.272 | 1 | - | - | - | - | - |
| | | Squared Activity (cm/sec, log-transformed) | 61.856 | 36.815 | 2.669 | 0.102 | 1 | - | - | - | - | - |
| | | Feeding (min) | 0.630 | 0.509 | 1.485 | 0.223 | 1 | - | - | - | - | - |
| | | **Random Effects** | | | | | | | | | | |
| | | $\sigma^2$ | 4212.98 | | | | | 4798.35 | | | | |
| | | $\tau_{00}$ | 0.00 Mother ID | | | | | 0.00 Mother ID | | | | |
| | | *N* | 3 Mother ID | | | | | 3 Mother ID | | | | |
| | | Observations | 24 | | | | | 24 | | | | |
| | | Marginal $R^2$ / Conditional $R^2$ | 0.127 / NA | | | | | 0.000 / NA | | | | |
| | | AIC | 255.774 | | | | | 270.377 | | | | |

**Supplementary Table 4.** Continued.

| Time interval | Response | Predictor | Full model | | | | | Final model | | | | |
|---|---|---|---|---|---|---|---|---|---|---|---|---|
| | | | **Estimate** | **SE** | **$\chi^2$** | **p** | **df** | **Estimate** | **SE** | **$\chi^2$** | **p** | **df** |
| Day 20 | Lifespan (days) | (Intercept) | 511.608 | 33.124 | - | - | - | 511.608 | 33.124 | - | - | - |
| | | Activity (cm/sec, log-transformed) | -23.029 | 23.576 | 0.941 | 0.332 | 1 | -23.029 | 23.576 | 0.941 | 0.332 | 1 |
| | | Squared Activity (cm/sec, log-transformed) | 109.916 | 41.370 | 6.397 | 0.011 | 1 | 109.916 | 41.370 | 6.397 | 0.011 | 1 |
| | | Feeding (min) | 1.524 | 0.383 | 12.947 | <0.001 | 1 | 1.524 | 0.383 | 12.947 | <0.001 | 1 |
| | | **Random Effects** | | | | | | | | | | |
| | | $\sigma^2$ | 3694.31 | | | | | 3694.31 | | | | |
| | | $\tau_{00}$ | 0.00 Mother ID | | | | | 0.00 Mother ID | | | | |
| | | *N* | 3 Mother ID | | | | | 3 Mother ID | | | | |
| | | Observations | 33 | | | | | 33 | | | | |
| | | Marginal $R^2$ / Conditional $R^2$ | 0.393 / NA | | | | | 0.393 / NA | | | | |
| | | AIC | 352.610 | | | | | 352.610 | | | | |

**Supplementary Table 4.** Continued.

| Time interval | Response | Predictor | Full model | | | | | Final model | | | | |
|---|---|---|---|---|---|---|---|---|---|---|---|---|
| | | | Estimate | SE | $\chi^2$ | p | df | Estimate | SE | $\chi^2$ | p | df |
| Day 21 | Lifespan (days) | (Intercept) | 565.420 | 38.303 | - | - | - | 653.394 | 13.498 | - | - | - |
| | | Activity (cm/sec, log-transformed) | -35.330 | 31.215 | 1.257 | 0.262 | 1 | - | - | - | - | - |
| | | Squared Activity (cm/sec, log-transformed) | 66.812 | 38.747 | 2.847 | 0.092 | 1 | - | - | - | - | - |
| | | Feeding (min) | 0.838 | 0.427 | 3.638 | 0.056 | 1 | - | - | - | - | - |
| | | **Random Effects** | | | | | | | | | | |
| | | $\sigma^2$ | 5060.03 | | | | | 6012.78 | | | | |
| | | $\tau_{00}$ | 0.00 Mother ID | | | | | 0.00 Mother ID | | | | |
| | | *N* | 3 Mother ID | | | | | 3 Mother ID | | | | |
| | | Observations | 33 | | | | | 33 | | | | |
| | | Marginal $R^2$ / Conditional $R^2$ | 0.163 / NA | | | | | 0.000 / NA | | | | |
| | | AIC | 362.101 | | | | | 379.746 | | | | |

**Supplementary Table 4.** Continued.

| Time interval | Response | Predictor | Full model | | | | | Final model | | | | |
|---|---|---|---|---|---|---|---|---|---|---|---|---|
| | | | Estimate | SE | $\chi^2$ | p | df | Estimate | SE | $\chi^2$ | p | df |
| Day 22 | Lifespan (days) | (Intercept) | 657.001 | 27.709 | - | - | - | 653.394 | 13.498 | - | - | - |
| | | Activity (cm/sec, log-transformed) | 4.531 | 35.247 | 0.017 | 0.898 | 1 | - | - | - | - | - |
| | | Squared Activity (cm/sec, log-transformed) | -16.014 | 24.844 | 0.413 | 0.521 | 1 | - | - | - | - | - |
| | | Feeding (min) | 0.181 | 0.479 | 0.143 | 0.706 | 1 | - | - | - | - | - |
| | | **Random Effects** | | | | | | | | | | |
| | | $\sigma^2$ | 5786.40 | | | | | 6012.78 | | | | |
| | | $\tau_{00}$ | 0.00 Mother ID | | | | | 0.00 Mother ID | | | | |
| | | *N* | 3 Mother ID | | | | | 3 Mother ID | | | | |
| | | Observations | 33 | | | | | 33 | | | | |
| | | Marginal $R^2$ / Conditional $R^2$ | 0.039 / NA | | | | | 0.000 / NA | | | | |
| | | AIC | 367.455 | | | | | 379.746 | | | | |

**Supplementary Table 4.** Continued.

| Time interval | Response | Predictor | Full model | | | | | Final model | | | | |
|---|---|---|---|---|---|---|---|---|---|---|---|---|
| | | | Estimate | SE | $\chi^2$ | p | df | Estimate | SE | $\chi^2$ | p | df |
| Day 23 | Lifespan (days) | (Intercept) | 665.557 | 29.807 | - | - | - | 653.394 | 13.498 | - | - | - |
| | | Activity (cm/sec, log-transformed) | 6.035 | 23.831 | 0.064 | 0.802 | 1 | - | - | - | - | - |
| | | Squared Activity (cm/sec, log-transformed) | -2.807 | 21.280 | 0.017 | 0.896 | 1 | - | - | - | - | - |
| | | Feeding (min) | -0.181 | 0.612 | 0.087 | 0.768 | 1 | - | - | - | - | - |
| | | **Random Effects** | | | | | | | | | | |
| | | $\sigma^2$ | 5962.27 | | | | | 6012.78 | | | | |
| | | $\tau_{00}$ | 0.00 Mother ID | | | | | 0.00 Mother ID | | | | |
| | | *N* | 3 Mother ID | | | | | 3 Mother ID | | | | |
| | | Observations | 33 | | | | | 33 | | | | |
| | | Marginal $R^2$ / Conditional $R^2$ | 0.009 / NA | | | | | 0.000 / NA | | | | |
| | | AIC | 368.484 | | | | | 379.746 | | | | |

**Supplementary Table 4.** Continued.

| Time interval | Response | Predictor | Full model | | | | | Final model | | | | |
|---|---|---|---|---|---|---|---|---|---|---|---|---|
| | | | Estimate | SE | $\chi^2$ | p | df | Estimate | SE | $\chi^2$ | p | df |
| Day 24 | Lifespan (days) | (Intercept) | 606.711 | 36.439 | - | - | - | 653.394 | 13.498 | - | - | - |
| | | Activity (cm/sec, log-transformed) | 4.246 | 31.955 | 0.018 | 0.894 | 1 | - | - | - | - | - |
| | | Squared Activity (cm/sec, log-transformed) | 23.373 | 51.820 | 0.203 | 0.653 | 1 | - | - | - | - | - |
| | | Feeding (min) | 0.757 | 0.550 | 1.842 | 0.175 | 1 | - | - | - | - | - |
| | | **Random Effects** | | | | | | | | | | |
| | | $\sigma^2$ | 5662.43 | | | | | 6012.78 | | | | |
| | | $\tau_{00}$ | 0.00 Mother ID | | | | | 0.00 Mother ID | | | | |
| | | *N* | 3 Mother ID | | | | | 3 Mother ID | | | | |
| | | Observations | 33 | | | | | 33 | | | | |
| | | Marginal $R^2$ / Conditional $R^2$ | 0.060 / NA | | | | | 0.000 / NA | | | | |
| | | AIC | 364.584 | | | | | 379.746 | | | | |

**Supplementary Table 5.** Continued.

| Time interval | Response | Predictor | Full model | | | | | Final model | | | | |
|---|---|---|---|---|---|---|---|---|---|---|---|---|
| | | | Estimate | SE | $\chi^2$ | p | df | Estimate | SE | $\chi^2$ | p | df |
| Day 25 | Lifespan (days) | (Intercept) | 632.017 | 35.048 | - | - | - | 653.394 | 13.498 | - | - | - |
| | | Activity (cm/sec, log-transformed) | 7.078 | 28.801 | 0.060 | 0.806 | 1 | - | - | - | - | - |
| | | Squared Activity (cm/sec, log-transformed) | -3.245 | 26.853 | 0.015 | 0.904 | 1 | - | - | - | - | - |
| | | Feeding (min) | 0.381 | 0.517 | 0.539 | 0.463 | 1 | - | - | - | - | - |
| | | **Random Effects** | | | | | | | | | | |
| | | $\sigma^2$ | 5915.29 | | | | | 6012.78 | | | | |
| | | $\tau_{00}$ | 0.00 Mother ID | | | | | 0.00 Mother ID | | | | |
| | | *N* | 3 Mother ID | | | | | 3 Mother ID | | | | |
| | | Observations | 33 | | | | | 33 | | | | |
| | | Marginal $R^2$ / Conditional $R^2$ | 0.017 / NA | | | | | 0.000 / NA | | | | |
| | | AIC | 367.815 | | | | | 379.746 | | | | |

**Supplementary Table 4.** Continued.

| Time interval | Response | Predictor | Full model | | | | | Final model | | | | |
|---|---|---|---|---|---|---|---|---|---|---|---|---|
| | | | Estimate | SE | $\chi^2$ | p | df | Estimate | SE | $\chi^2$ | p | df |
| Day 26 | Lifespan (days) | (Intercept) | 651.777 | 30.951 | - | - | - | 653.394 | 13.498 | - | - | - |
| | | Activity (cm/sec, log-transformed) | -28.238 | 33.822 | 0.691 | 0.406 | 1 | - | - | - | - | - |
| | | Squared Activity (cm/sec, log-transformed) | 24.436 | 24.931 | 0.947 | 0.331 | 1 | - | - | - | - | - |
| | | Feeding (min) | -0.334 | 0.523 | 0.406 | 0.524 | 1 | - | - | - | - | - |
| | | **Random Effects** | | | | | | | | | | |
| | | $\sigma^2$ | 5822.91 | | | | | 6012.78 | | | | |
| | | $\tau_{00}$ | 0.00 Mother ID | | | | | 0.00 Mother ID | | | | |
| | | *N* | 3 Mother ID | | | | | 3 Mother ID | | | | |
| | | Observations | 33 | | | | | 33 | | | | |
| | | Marginal $R^2$ / Conditional $R^2$ | 0.033 / NA | | | | | 0.000 / NA | | | | |
| | | AIC | 367.654 | | | | | 379.746 | | | | |

**Supplementary Table 4.** Continued.

| **Time interval** | **Response** | **Predictor** | **Full model** | | | | | **Final model** | | | | |
|---|---|---|---|---|---|---|---|---|---|---|---|---|
| | | | **Estimate** | **SE** | **$\chi^2$** | **p** | **df** | **Estimate** | **SE** | **$\chi^2$** | **p** | **df** |
| Day 27 | Lifespan (days) | (Intercept) | 618.767 | 29.116 | - | - | - | 653.394 | 13.498 | - | - | - |
| | | Activity (cm/sec, log-transformed) | 22.980 | 25.745 | 0.787 | 0.375 | 1 | - | - | - | - | - |
| | | Squared Activity (cm/sec, log-transformed) | 9.016 | 24.318 | 0.137 | 0.711 | 1 | - | - | - | - | - |
| | | Feeding (min) | 0.729 | 0.497 | 2.080 | 0.149 | 1 | - | - | - | - | - |
| | | **Random Effects** | | | | | | | | | | |
| | | $\sigma^2$ | 5508.62 | | | | | 6012.78 | | | | |
| | | $\tau_{00}$ | 0.00 Mother ID | | | | | 0.00 Mother ID | | | | |
| | | *N* | 3 Mother ID | | | | | 3 Mother ID | | | | |
| | | Observations | 33 | | | | | 33 | | | | |
| | | Marginal $R^2$ / Conditional $R^2$ | 0.086 / NA | | | | | 0.000 / NA | | | | |
| | | AIC | 366.061 | | | | | 379.746 | | | | |

**Supplementary Table 4.** Continued.

| Time interval | Response | Predictor | Full model | | | | | Final model | | | | |
|---|---|---|---|---|---|---|---|---|---|---|---|---|
| | | | Estimate | SE | $\chi^2$ | p | df | Estimate | SE | $\chi^2$ | p | df |
| Day 28 | Lifespan (days) | (Intercept) | 611.281 | 32.109 | - | - | - | 656.759 | 14.879 | - | - | - |
| | | Activity (cm/sec, log-transformed) | 12.477 | 31.379 | 0.158 | 0.691 | 1 | - | - | - | - | - |
| | | Squared Activity (cm/sec, log-transformed) | 14.738 | 28.004 | 0.276 | 0.611 | 1 | - | - | - | - | - |
| | | Feeding (min) | 0.835 | 0.602 | 1.862 | 0.172 | 1 | - | - | - | - | - |
| | | **Random Effects** | | | | | | | | | | |
| | | $\sigma^2$ | 5785.06 | | | | | 6419.98 | | | | |
| | | $\tau_{00}$ | 0.00 Mother ID | | | | | 0.00 Mother ID | | | | |
| | | *N* | 2 Mother ID | | | | | 2 Mother ID | | | | |
| | | Observations | 29 | | | | | 29 | | | | |
| | | Marginal $R^2$ / Conditional $R^2$ | 0.102 / NA | | | | | 0.000 / NA | | | | |
| | | AIC | 320.821 | | | | | 335.291 | | | | |

#### Supplementary Note 4: Indirect feeding-lifespan link via body size

**Supplementary Table 5. Summary of linear mixed-effects models testing whether early-life feeding behavior predicts maximum body size.** Maximum body size was estimated from individual van Bertalanffy growth curves. Models were first fitted using feeding behavior averaged across the entire first four weeks of life, and subsequently refitted with feeding behavior calculated separately for each of the four weeks to assess the temporal development of this relationship.

| Time interval | Response | Predictor | Full model | | | | | Final model | | | | |
|---|---|---|---|---|---|---|---|---|---|---|---|---|
| | | | Estimate | SE | $\chi^2$ | p | df | Estimate | SE | $\chi^2$ | p | df |
| Week 1-4 | Maximum body size | (Intercept) | 4.966 | 0.203 | - | - | - | 4.966 | 0.203 | - | - | - |
| | | Feeding (min) | 0.008 | 0.004 | 4.467 | 0.035 | 1 | 0.008 | 0.004 | 4.467 | 0.035 | 1 |
| | | **Random Effects** | | | | | | | | | | |
| | | $\sigma^2$ | 0.09 | | | | | 0.09 | | | | |
| | | $\tau_{00}$ | 0.03 Mother ID | | | | | 0.03 Mother ID | | | | |
| | | ICC | 0.21 | | | | | 0.21 | | | | |
| | | *N* | 3 Mother ID | | | | | 3 Mother ID | | | | |
| | | Observations | 33 | | | | | 33 | | | | |
| | | Marginal $R^2$ / Conditional $R^2$ | 0.112 / 0.302 | | | | | 0.112 / 0.302 | | | | |
| | | AIC | 38.840 | | | | | 38.840 | | | | |

**Supplementary Table 5.** Continued.

| Time interval | Response | Predictor | Full model | | | | | Final model | | | | |
|---|---|---|---|---|---|---|---|---|---|---|---|---|
| | | | **Estimate** | **SE** | **$\chi^2$** | **p** | **df** | **Estimate** | **SE** | **$\chi^2$** | **p** | **df** |
| Week 1 | Maximum body size | (Intercept) | 5.218 | 0.137 | - | - | - | 5.341 | 0.122 | - | - | - |
| | | Feeding (min) | 0.005 | 0.003 | 3.133 | 0.077 | 1 | - | - | - | - | - |
| | | **Random Effects** | | | | | | | | | | |
| | | $\sigma^2$ | 0.09 | | | | | 0.10 | | | | |
| | | $\tau_{00}$ | 0.03 Mother ID | | | | | 0.03 Mother ID | | | | |
| | | ICC | 0.25 | | | | | 0.24 | | | | |
| | | *N* | 3 Mother ID | | | | | 3 Mother ID | | | | |
| | | Observations | 33 | | | | | 33 | | | | |
| | | Marginal $R^2$ / Conditional $R^2$ | 0.075 / 0.304 | | | | | 0.000 / 0.239 | | | | |
| | | AIC | 40.472 | | | | | 31.739 | | | | |

**Supplementary Table 5.** Continued.

| Time interval | Response | Predictor | Full model | | | | | Final model | | | | |
|---|---|---|---|---|---|---|---|---|---|---|---|---|
| | | | Estimate | SE | $\chi^2$ | p | df | Estimate | SE | $\chi^2$ | p | df |
| Week 2 | Maximum body size | (Intercept) | 5.290 | 0.178 | - | - | - | 5.341 | 0.122 | - | - | - |
| | | Feeding (min) | 0.001 | 0.003 | 0.143 | 0.705 | 1 | - | - | - | - | - |
| | | **Random Effects** | | | | | | | | | | |
| | | $\sigma^2$ | 0.10 | | | | | 0.10 | | | | |
| | | $\tau_{00}$ | 0.03 Mother ID | | | | | 0.03 Mother ID | | | | |
| | | ICC | 0.22 | | | | | 0.24 | | | | |
| | | *N* | 3 Mother ID | | | | | 3 Mother ID | | | | |
| | | Observations | 33 | | | | | 33 | | | | |
| | | Marginal $R^2$ / Conditional $R^2$ | 0.004 / 0.226 | | | | | 0.000 / 0.239 | | | | |
| | | AIC | 43.692 | | | | | 31.739 | | | | |

**Supplementary Table 5.** Continued.

| Time interval | Response | Predictor | Full model | | | | | Final model | | | | |
|---|---|---|---|---|---|---|---|---|---|---|---|---|
| | | | Estimate | SE | $\chi^2$ | p | df | Estimate | SE | $\chi^2$ | p | df |
| Week 3 | Maximum body size | (Intercept) | 4.791 | 0.210 | - | - | - | 4.791 | 0.210 | - | - | - |
| | | Feeding (min) | 0.009 | 0.003 | 7.431 | 0.006 | 1 | 0.009 | 0.003 | 7.431 | 0.006 | 1 |
| | | **Random Effects** | | | | | | | | | | |
| | | $\sigma^2$ | 0.09 | | | | | 0.09 | | | | |
| | | $\tau_{00}$ | 0.02 Mother ID | | | | | 0.02 Mother ID | | | | |
| | | ICC | 0.16 | | | | | 0.16 | | | | |
| | | *N* | 3 Mother ID | | | | | 3 Mother ID | | | | |
| | | Observations | 33 | | | | | 33 | | | | |
| | | Marginal $R^2$ / Conditional $R^2$ | 0.201 / 0.331 | | | | | 0.201 / 0.331 | | | | |
| | | AIC | 36.478 | | | | | 36.478 | | | | |

**Supplementary Table 5.** Continued.

| Time interval | Response | Predictor | Full model | | | | | Final model | | | | |
|---|---|---|---|---|---|---|---|---|---|---|---|---|
| | | | Estimate | SE | $\chi^2$ | p | df | Estimate | SE | $\chi^2$ | p | df |
| Week 4 | Maximum body size | (Intercept) | 5.081 | 0.179 | - | - | - | 5.341 | 0.122 | - | - | - |
| | | Feeding (min) | 0.005 | 0.003 | 1.979 | 0.057 | 1 | - | - | - | - | - |
| | | **Random Effects** | | | | | | | | | | |
| | | $\sigma^2$ | 0.09 | | | | | 0.10 | | | | |
| | | $\tau_{00}$ | 0.03 Mother ID | | | | | 0.03 Mother ID | | | | |
| | | ICC | 0.27 | | | | | 0.24 | | | | |
| | | *N* | 3 Mother ID | | | | | 3 Mother ID | | | | |
| | | Observations | 33 | | | | | 33 | | | | |
| | | Marginal $R^2$ / Conditional $R^2$ | 0.083 / 0.331 | | | | | 0.000 / 0.239 | | | | |
| | | AIC | 40.128 | | | | | 31.739 | | | | |

**Supplementary Table 6. Summary of the linear mixed-effects model testing whether maximum body size predicts lifespan.** Maximum body size was estimated for each individual from van Bertalanffy growth curves and estimated as a predictor of lifespan. The model assesses whether size-related variation in growth trajectories translates into differences in longevity.

| ***Response*** | ***Predictors*** | **Full and final model** | | | | |
|---|---|---|---|---|---|---|
| | | ***Estimate*** | ***SE*** | **$\chi^2$** | ***p*** | ***df*** |
| Lifespan (days) | (Intercept) | 260.093 | 179.152 | - | - | - |
| | Maximum body size | 73.412 | 33.357 | 4.519 | 0.034 | 1 |
| | **Random Effects** | | | | | |
| | $\sigma^2$ | 5243.22 | | | | |
| | $\tau_{00}$ | 0.00 Mother ID | | | | |
| | *N* | 3 Mother ID | | | | |
| | Observations | 33 | | | | |
| | Marginal $R^2$ / Conditional $R^2$ | 0.131 / NA | | | | |
| | AIC | 368.464 | | | | |

#### Supplementary Note 5: Robustness with respect to size at birth

**Supplementary Table 7. Summary of linear mixed-effects models testing whether early-life behavior predicts lifespan while controlling for size at birth.** These models mirror the primary analysis presented in Supplementary Table 1, with the addition of offspring size at birth as a covariate to assess whether the activity–lifespan relationship is mediated by body size at birth. As in the primary analysis, behavior was first averaged across the entire first four weeks of life and then analyzed at finer temporal resolution by fitting separate models for each of the four weeks individually, allowing assessment of developmental dynamics.

| Time interval | Response | Predictor | Full model | | | | | Final model | | | | |
|---|---|---|---|---|---|---|---|---|---|---|---|---|
| | | | Estimate | SE | $\chi^2$ | p | df | Estimate | SE | $\chi^2$ | p | df |
| Week 1-4 | Lifespan (days) | (Intercept) | 571.277 | 83.389 | - | - | - | 636.824 | 14.390 | - | - | - |
| | | Activity (cm/sec, log-transformed) | -138.387 | 37.499 | 11.402 | 0.001 | 1 | -142.148 | 36.375 | -3.908 | 0.001 | 1 |
| | | Squared Activity (cm/sec, log-transformed) | 132.546 | 43.450 | 8.198 | 0.005 | 1 | 129.667 | 44.109 | 2.940 | 0.007 | 1 |
| | | Feeding (min) | 0.736 | 0.836 | 0.768 | 0.381 | 1 | - | - | - | - | - |
| | | Size at birth (cm) | 22.044 | 65.877 | 0.112 | 0.738 | 1 | - | - | - | - | - |
| | | **Random Effects** | | | | | | | | | | |
| | | $\sigma^2$ | 3957.28 | | | | | 4098.82 | | | | |
| | | $\tau_{00}$ | 0.00 Mother ID | | | | | 0.00 Mother ID | | | | |
| | | *N* | 3 Mother ID | | | | | 3 Mother ID | | | | |
| | | Observations | 33 | | | | | 33 | | | | |
| | | Marginal $R^2$ / Conditional $R^2$ | 0.349 / NA | | | | | 0.325 / NA | | | | |
| | | AIC | 345.075 | | | | | 353.981 | | | | |

**Supplementary Table 7.** Continued.

| ***Time interval*** | ***Response*** | ***Predictor*** | **Full model** | | | | | **Final model** | | | | |
|---|---|---|---|---|---|---|---|---|---|---|---|---|
| | | | ***Estimate*** | ***SE*** | $\chi^2$ | ***p*** | ***df*** | ***Estimate*** | ***SE*** | $\chi^2$ | ***p*** | ***df*** |
| Week 1 | Lifespan (days) | (Intercept) | 684.488 | 83.781 | - | - | - | 699.570 | 17.412 | - | - | - |
| | | Activity (cm/sec, log-transformed) | -67.915 | 60.790 | 1.225 | 0.268 | 1 | -64.740 | 18.331 | -3.532 | 0.001 | 1 |
| | | Squared Activity (cm/sec, log-transformed) | 9.969 | 36.923 | 0.073 | 0.787 | 1 | - | - | - | - | - |
| | | Feeding (min) | 0.747 | 0.715 | 1.073 | 0.300 | 1 | - | - | - | - | - |
| | | Size at birth (cm) | -8.124 | 63.163 | 0.017 | 0.898 | 1 | - | - | - | - | - |
| | | **Random Effects** | | | | | | | | | | |
| | | $\sigma^2$ | 4202.19 | | | | | 4363.52 | | | | |
| | | $\tau_{00}$ | 0.00 Mother ID | | | | | 0.00 Mother ID | | | | |
| | | *N* | 3 Mother ID | | | | | 3 Mother ID | | | | |
| | | Observations | 33 | | | | | 33 | | | | |
| | | Marginal $R^2$ / Conditional $R^2$ | 0.308 / NA | | | | | 0.280 / NA | | | | |
| | | AIC | 347.986 | | | | | 363.785 | | | | |

**Supplementary Table 7.** Continued.

| Time interval | Response | Predictor | Full model | | | | | Final model | | | | |
|---|---|---|---|---|---|---|---|---|---|---|---|---|
| | | | **Estimate** | **SE** | **$\chi^2$** | **p** | **df** | **Estimate** | **SE** | **$\chi^2$** | **p** | **df** |
| Week 2 | Lifespan (days) | (Intercept) | 621.423 | 86.420 | - | - | - | 649.157 | 12.598 | - | - | - |
| | | Activity (cm/sec, log-transformed) | -96.034 | 29.811 | 9.023 | 0.003 | 1 | -52.860 | 22.221 | -2.379 | 0.024 | 1 |
| | | Squared Activity (cm/sec, log-transformed) | 60.894 | 32.259 | 3.384 | 0.067 | 1 | - | - | - | - | - |
| | | Feeding (min) | -0.463 | 0.577 | 0.638 | 0.424 | 1 | - | - | - | - | - |
| | | Size at birth (cm) | 22.653 | 66.828 | 0.115 | 0.735 | 1 | - | - | - | - | - |
| | | **Random Effects** | | | | | | | | | | |
| | | $\sigma^2$ | 4551.65 | | | | | 5132.66 | | | | |
| | | $\tau_{00}$ | 0.00 Mother ID | | | | | 0.00 Mother ID | | | | |
| | | *N* | 3 Mother ID | | | | | 3 Mother ID | | | | |
| | | Observations | 33 | | | | | 33 | | | | |
| | | Marginal $R^2$ / Conditional $R^2$ | 0.249 / NA | | | | | 0.150 / NA | | | | |
| | | AIC | 350.832 | | | | | 368.595 | | | | |

**Supplementary Table 7.** Continued.

| Time interval | Response | Predictor | Full model | | | | | Final model | | | | |
|---|---|---|---|---|---|---|---|---|---|---|---|---|
| | | | **Estimate** | **SE** | **$\chi^2$** | **p** | **df** | **Estimate** | **SE** | **$\chi^2$** | **p** | **df** |
| Week 3 | Lifespan (days) | (Intercept) | 502.134 | 102.726 | - | - | - | 588.510 | 27.139 | - | - | - |
| | | Activity (cm/sec, log-transformed) | -72.272 | 31.174 | 4.981 | 0.026 | 1 | -65.485 | 30.116 | 4.419 | 0.036 | 1 |
| | | Squared Activity (cm/sec, log-transformed) | 117.116 | 37.647 | 8.4487 | 0.004 | 1 | 103.906 | 38.533 | 6.571 | 0.010 | 1 |
| | | Feeding (min) | 1.380 | 0.720 | 3.488 | 0.062 | 1 | - | - | - | - | - |
| | | Size at birth (cm) | -4.433 | 73.335 | 0.004 | 0.952 | 1 | - | - | - | - | - |
| | | **Random Effects** | | | | | | | | | | |
| | | $\sigma^2$ | 4391.42 | | | | | 4921.67 | | | | |
| | | $\tau_{00}$ | 0.00 Mother ID | | | | | 0.00 Mother ID | | | | |
| | | *N* | 3 Mother ID | | | | | 3 Mother ID | | | | |
| | | Observations | 33 | | | | | 33 | | | | |
| | | Marginal $R^2$ / Conditional $R^2$ | 0.276 / NA | | | | | 0.186 / NA | | | | |
| | | AIC | 349.080 | | | | | 360.268 | | | | |

**Supplementary Table 7.** Continued.

| Time interval | Response | Predictor | Full model | | | | | Final model | | | | |
|---|---|---|---|---|---|---|---|---|---|---|---|---|
| | | | Estimate | SE | $\chi^2$ | p | df | Estimate | SE | $\chi^2$ | p | df |
| Week 4 | Lifespan (days) | (Intercept) | 663.117 | 116.794 | - | - | - | 653.394 | 13.498 | - | - | - |
| | | Activity (cm/sec, log-transformed) | 15.058 | 38.563 | 0.152 | 0.697 | 1 | - | - | - | - | - |
| | | Squared Activity (cm/sec, log-transformed) | 12.911 | 45.497 | 0.080 | 0.777 | 1 | - | - | - | - | - |
| | | Feeding (min) | 0.577 | 0.723 | 0.631 | 0.427 | 1 | - | - | - | - | - |
| | | Size at birth (cm) | -30.826 | 83.470 | 0.136 | 0.712 | 1 | - | - | - | - | - |
| | | **Random Effects** | | | | | | | | | | |
| | | $\sigma^2$ | 5802.22 | | | | | 6012.78 | | | | |
| | | $\tau_{00}$ | 0.00 Mother ID | | | | | 0.00 Mother ID | | | | |
| | | *N* | 3 Mother ID | | | | | 3 Mother ID | | | | |
| | | Observations | 33 | | | | | 33 | | | | |
| | | Marginal $R^2$ / Conditional $R^2$ | 0.036 / NA | | | | | 0.000 / NA | | | | |
| | | AIC | 356.765 | | | | | 379.746 | | | | |

#### Supplementary Note 6: Robustness with respect to the removal of five outlier individuals

**Supplementary Table 8. Robustness of early-life behavior–lifespan relationships to outlier inclusion.** Summary of linear mixed-effects models testing whether early-life activity and feeding behavior predict lifespan, using the same model structure as in the primary analysis (Supplementary Table 1) but including the five previously excluded outlier individuals. As in the primary analysis, activity and feeding behavior were averaged either across the full four-week observation period or separately for each of the first four weeks of life, resulting in five models in total.

| Time interval | Response | Predictor | Full model | | | | | Final model | | | | |
|---|---|---|---|---|---|---|---|---|---|---|---|---|
| | | | **Estimate** | **SE** | **$\chi^2$** | **p** | **df** | **Estimate** | **SE** | **$\chi^2$** | **p** | **df** |
| Week 1-4 | Lifespan (days) | (Intercept) | 415.702 | 88.320 | - | - | - | 563.352 | 31.648 | - | - | - |
| | | Activity (cm/sec, log-transformed) | -152.992 | 82.994 | 3.255 | 0.071 | 1 | -188.647 | 83.835 | 4.753 | 0.029 | 1 |
| | | Squared Activity (cm/sec, log-transformed) | 219.195 | 98.021 | 4.698 | 0.030 | 1 | 224.348 | 101.984 | 4.555 | 0.033 | 1 |
| | | Feeding (min) | 3.054 | 1.715 | 3.045 | 0.081 | 1 | - | - | - | - | - |
| | | **Random Effects** | | | | | | | | | | |
| | | $\sigma^2$ | 21571.35 | | | | | 23371.16 | | | | |
| | | $\tau_{00}$ | 0.00 Mother ID | | | | | 0.00 Mother ID | | | | |
| | | *N* | 3 Mother ID | | | | | 3 Mother ID | | | | |
| | | Observations | 38 | | | | | 38 | | | | |
| | | Marginal $R^2$ / Conditional $R^2$ | 0.197 / NA | | | | | 0.129 / NA | | | | |
| | | AIC | 467.186 | | | | | 471.004 | | | | |

**Supplementary Table 8.** Continued.

| Time interval | Response | Predictor | Full model | | | | | Final model | | | | |
|---|---|---|---|---|---|---|---|---|---|---|---|---|
| | | | Estimate | SE | $\chi^2$ | p | df | Estimate | SE | $\chi^2$ | p | df |
| Week 1 | Lifespan (days) | (Intercept) | 573.281 | 68.837 | - | - | - | 516.785 | 39.968 | - | - | - |
| | | Activity (cm/sec, log-transformed) | -206.294 | 133.565 | 2.314 | 0.128 | 1 | - | - | - | - | - |
| | | Squared Activity (cm/sec, log-transformed) | 124.250 | 80.016 | 2.338 | 0.126 | 1 | - | - | - | - | - |
| | | Feeding (min) | 2.668 | 1.508 | 3.011 | 0.083 | 1 | 3.388 | 1.327 | 6.019 | 0.014 | 1 |
| | | **Random Effects** | | | | | | | | | | |
| | | $\sigma^2$ | 21425.74 | | | | | 22819.35 | | | | |
| | | $\tau_{00}$ | 0.00 Mother ID | | | | | 0.00 Mother ID | | | | |
| | | *N* | 3 Mother ID | | | | | 3 Mother ID | | | | |
| | | Observations | 38 | | | | | 38 | | | | |
| | | Marginal $R^2$ / Conditional $R^2$ | 0.203 / NA | | | | | 0.150 / NA | | | | |
| | | AIC | 467.998 | | | | | 486.491 | | | | |

**Supplementary Table 8.** Continued.

| Time interval | Response | Predictor | Full model | | | | | Final model | | | | |
|---|---|---|---|---|---|---|---|---|---|---|---|---|
| | | | Estimate | SE | $\chi^2$ | p | df | Estimate | SE | $\chi^2$ | p | df |
| Week 2 | Lifespan (days) | (Intercept) | 509.568 | 68.091 | - | - | - | 597.421 | 26.525 | - | - | - |
| | | Activity (cm/sec, log-transformed) | -54.911 | 64.278 | 0.723 | 0.395 | 1 | - | - | - | - | - |
| | | Squared Activity (cm/sec, log-transformed) | 76.187 | 71.943 | 1.105 | 0.293 | 1 | - | - | - | - | - |
| | | Feeding (min) | 1.157 | 1.195 | 0.926 | 0.336 | 1 | - | - | - | - | - |
| | | **Random Effects** | | | | | | | | | | |
| | | $\sigma^2$ | 24869.57 | | | | | 26735.98 | | | | |
| | | $\tau_{00}$ | 0.00 Mother ID | | | | | 0.00 Mother ID | | | | |
| | | *N* | 3 Mother ID | | | | | 3 Mother ID | | | | |
| | | Observations | 38 | | | | | 38 | | | | |
| | | Marginal $R^2$ / Conditional $R^2$ | 0.072 / NA | | | | | 0.000 / NA | | | | |
| | | AIC | 473.825 | | | | | 492.795 | | | | |

**Supplementary Table 8.** Continued.

| Time interval | Response | Predictor | Full model | | | | | Final model | | | | |
|---|---|---|---|---|---|---|---|---|---|---|---|---|
| | | | Estimate | SE | $\chi^2$ | p | df | Estimate | SE | $\chi^2$ | p | df |
| Week 3 | Lifespan (days) | (Intercept) | 350.293 | 101.977 | - | - | - | 597.421 | 26.525 | - | - | - |
| | | Activity (cm/sec, log-transformed) | -81.072 | 57.257 | 1.954 | 0.162 | 1 | - | - | - | - | - |
| | | Squared Activity (cm/sec, log-transformed) | 159.461 | 76.025 | 4.163 | 0.041 | 1 | - | - | - | - | - |
| | | Feeding (min) | 2.602 | 1.433 | 3.164 | 0.075 | 1 | - | - | - | - | - |
| | | **Random Effects** | | | | | | | | | | |
| | | $\sigma^2$ | 22440.70 | | | | | 26735.98 | | | | |
| | | $\tau_{00}$ | 0.00 Mother ID | | | | | 0.00 Mother ID | | | | |
| | | *N* | 3 Mother ID | | | | | 3 Mother ID | | | | |
| | | Observations | 38 | | | | | 38 | | | | |
| | | Marginal $R^2$ / Conditional $R^2$ | 0.164 / NA | | | | | 0.000 / NA | | | | |
| | | AIC | 469.842 | | | | | 492.795 | | | | |

**Supplementary Table 8.** Continued.

| Time interval | Response | Predictor | Full model | | | | | Final model | | | | |
|---|---|---|---|---|---|---|---|---|---|---|---|---|
| | | | Estimate | SE | $\chi^2$ | p | df | Estimate | SE | $\chi^2$ | p | df |
| Week 4 | Lifespan (days) | (Intercept) | 546.615 | 73.642 | - | - | - | 597.421 | 26.525 | - | - | - |
| | | Activity (cm/sec, log-transformed) | -25.660 | 65.949 | 0.151 | 0.698 | 1 | - | - | - | - | - |
| | | Squared Activity (cm/sec, log-transformed) | 99.613 | 90.056 | 1.204 | 0.273 | 1 | - | - | - | - | - |
| | | Feeding (min) | 0.155 | 1.411 | 0.012 | 0.913 | 1 | - | - | - | - | - |
| | | **Random Effects** | | | | | | | | | | |
| | | $\sigma^2$ | 25725.31 | | | | | 26735.98 | | | | |
| | | $\tau_{00}$ | 0.00 Mother ID | | | | | 0.00 Mother ID | | | | |
| | | *N* | 3 Mother ID | | | | | 3 Mother ID | | | | |
| | | Observations | 38 | | | | | 38 | | | | |
| | | Marginal $R^2$ / Conditional $R^2$ | 0.039 / NA | | | | | 0.000 / NA | | | | |
| | | AIC | 474.212 | | | | | 492.795 | | | | |

#### Supplementary Note 7: Lifespan in relation to additional behavioral and reproductive measures

**Supplementary Table 9. Linear mixed-effects model summaries testing whether early-life behavior predicts behavior in standard behavioral assays.** Three models assess whether activity and feeding behavior averaged over the first four weeks of life predict (i) activity in a novel tank, (ii) response to a novel object, and (iii) sociability. Behavioral assays were conducted directly after the early-life observation period. See Supplementary Note 9 for a detailed description of the experimental procedure of the standard behavioral assays.

| **Response** | **Predictor** | **Full model** | | | | | **Final model** | | | | |
|---|---|---|---|---|---|---|---|---|---|---|---|
| | | **Estimate** | **SE** | **$\chi^2$** | **p** | **df** | **Estimate** | **SE** | **$\chi^2$** | **p** | **df** |
| Activity in a new tank (cm/sec, log-transformed) | (Intercept) | 0.061 | 0.256 | - | - | - | 0.200 | 0.071 | - | - | - |
| | Activity (cm/sec, log-transformed) | 0.483 | 0.142 | 9.963 | 0.002 | 1 | 0.453 | 0.132 | 10.092 | 0.001 | 1 |
| | Feeding (min) | 0.003 | 0.005 | 0.320 | 0.572 | 1 | - | - | - | - | - |
| | **Random Effects** | | | | | | | | | | |
| | $\sigma^2$ | 0.15 | | | | | 0.16 | | | | |
| | $\tau_{00}$ | 0.00 Mother ID | | | | | 0.00 Mother ID | | | | |
| | *N* | 3 Mother ID | | | | | 3 Mother ID | | | | |
| | Observations | 33 | | | | | 33 | | | | |
| | Marginal $R^2$ / Conditional $R^2$ | 0.277 / NA | | | | | 0.269 / NA | | | | |
| | AIC | 73.996 | | | | | 63.594 | | | | |

**Supplementary Table 9.** Continued.

| Response | Predictor | Full model | | | | | Final model | | | | |
|---|---|---|---|---|---|---|---|---|---|---|---|
| | | Estimate | SE | $\chi^2$ | p | df | Estimate | SE | $\chi^2$ | p | df |
| Average distance to novel object (cm) | (Intercept) | 29.629 | 2.316 | - | - | - | 26.245 | 0.842 | - | - | - |
| | Activity (cm/sec, log-transformed) | -1.443 | 1.280 | 1.216 | 0.270 | 1 | - | - | - | - | - |
| | Feeding (min) | -0.070 | 0.045 | 2.019 | 0.155 | 1 | - | - | - | - | - |
| | **Random Effects** | | | | | | | | | | |
| | $\sigma^2$ | 12.54 | | | | | 12.82 | | | | |
| | $\tau_{00}$ | 0.00 Mother ID | | | | | 0.83 Mother ID | | | | |
| | ICC | - | | | | | 0.06 | | | | |
| | *N* | 3 Mother ID | | | | | 3 Mother ID | | | | |
| | Observations | 33 | | | | | 33 | | | | |
| | Marginal $R^2$ / Conditional $R^2$ | 0.080 / NA | | | | | 0.000 / 0.061 | | | | |
| | AIC | 188.295 | | | | | 183.885 | | | | |

**Supplementary Table 9.** Continued.

| Response | Predictor | Full model | | | | | Final model | | | | |
|---|---|---|---|---|---|---|---|---|---|---|---|
| | | Estimate | SE | $\chi^2$ | p | df | Estimate | SE | $\chi^2$ | p | df |
| Sociability (arcsine square root transformed) | (Intercept) | 1.416 | 0.057 | - | - | - | 1.352 | 0.016 | - | - | - |
| | Activity (cm/sec, log-transformed) | -0.067 | 0.031 | 4.289 | 0.038 | 1 | - | - | - | - | - |
| | Feeding (min) | -0.001 | 0.001 | 1.003 | 0.317 | 1 | - | - | - | - | - |
| | **Random Effects** | | | | | | | | | | |
| | $\sigma^2$ | 0.01 | | | | | 0.01 | | | | |
| | $\tau_{00}$ | 0.00 Mother ID | | | | | 0.00 Mother ID | | | | |
| | *N* | 3 Mother ID | | | | | 3 Mother ID | | | | |
| | Observations | 33 | | | | | 33 | | | | |
| | Marginal $R^2$ / Conditional $R^2$ | 0.126 / NA | | | | | 0.000 / NA | | | | |

**Supplementary Table 10. Summary of a linear mixed-effects model testing whether behavior expressed during standardized behavioral assays predicts lifespan.** The model assesses whether activity in a novel tank, response to a novel object, and sociability were predictive of lifespan. Behavioral assays were conducted directly after the early-life observation period. See Supplementary Note 9 for a detailed description of the experimental procedure of the standard behavioral assays.

| Response | Predictor | Full model | | | | | Final model | | | | |
|---|---|---|---|---|---|---|---|---|---|---|---|
| | | Estimate | SE | $\chi^2$ | p | df | Estimate | SE | $\chi^2$ | p | df |
| Lifespan (days) | (Intercept) | 711.290 | 210.685 | - | - | - | 653.394 | 13.498 | - | - | - |
| | Activity in a new tank (cm/sec, log-transformed) | -7.789 | 30.151 | 0.067 | 0.796 | 1 | - | - | - | - | - |
| | Averge distance to novel object (cm) | -6.104 | 3.687 | 2.633 | 0.105 | 1 | - | - | - | - | - |
| | Sociability (arcsine square root transformed) | 76.349 | 150.099 | 0.258 | 0.612 | 1 | - | - | - | - | - |
| | **Random Effects** | | | | | | | | | | |
| | $\sigma^2$ | 5545.59 | | | | | 6012.78 | | | | |
| | $\tau_{00}$ | 0.00 Mother ID | | | | | 0.00 Mother ID | | | | |
| | *N* | 3 Mother ID | | | | | 3 Mother ID | | | | |
| | Observations | 33 | | | | | 33 | | | | |
| | Marginal $R^2$ / Conditional $R^2$ | 0.080 / NA | | | | | 0.000 / NA | | | | |
| | AIC | 358.129 | | | | | 379.746 | | | | |

**Supplementary Table 11. Linear mixed-effects model summaries testing whether early-life behavior predicts later-life behavior.** Two models are shown: one testing whether early-life activity and feeding predict activity measured later in life, and a second testing whether the same early-life behavioral measures predict later-life feeding behavior. Early-life behavior was averaged across the first four weeks of life.

| Response | Predictor | Full model | | | | | Final model | | | | |
|---|---|---|---|---|---|---|---|---|---|---|---|
| | | Estimate | SE | $\chi^2$ | p | df | Estimate | SE | $\chi^2$ | p | df |
| Later-in-life activity (cm/sec, log-transformed) | (Intercept) | -7.716 | 0.796 | - | - | - | -8.047 | 0.700 | - | - | - |
| | Activity (cm/sec, log-transformed) | -0.366 | 0.434 | 0.703 | 0.402 | 1 | - | - | - | - | - |
| | Feeding (min) | -0.036 | 0.015 | 5.341 | 0.021 | 1 | -0.031 | 0.014 | 4.711 | 0.030 | 1 |
| | **Random Effects** | | | | | | | | | | |
| | $\sigma^2$ | 1.31 | | | | | 1.34 | | | | |
| | $\tau_{00}$ | 0.00 Mother ID | | | | | 0.00 Mother ID | | | | |
| | *N* | 3 Mother ID | | | | | 3 Mother ID | | | | |
| | Observations | 32 | | | | | 32 | | | | |
| | Marginal $R^2$ / Conditional $R^2$ | 0.160 / NA | | | | | 0.141 / NA | | | | |
| | AIC | 117.188 | | | | | 116.096 | | | | |

**Supplementary Table 11.** Continued.

| Response | Predictor | Full model | | | | | Final model | | | | |
|---|---|---|---|---|---|---|---|---|---|---|---|
| | | Estimate | SE | $\chi^2$ | p | df | Estimate | SE | $\chi^2$ | p | df |
| Later-in-life feeding (min) | (Intercept) | 1.340 | 15.636 | - | - | - | 32.206 | 4.228 | - | - | - |
| | Feeding (min) | 0.611 | 0.297 | 3.992 | 0.046 | 1 | - | - | - | - | - |
| | Activity (cm/sec, log-transformed) | 6.453 | 8.522 | 0.568 | 0.451 | 1 | - | - | - | - | - |
| | **Random Effects** | | | | | | | | | | |
| | $\sigma^2$ | 504.68 | | | | | 572.10 | | | | |
| | $\tau_{00}$ | 0.00 Mother ID | | | | | 0.00 Mother ID | | | | |
| | *N* | 3 Mother ID | | | | | 3 Mother ID | | | | |
| | Observations | 32 | | | | | 32 | | | | |
| | Marginal $R^2$ / Conditional $R^2$ | 0.121 / NA | | | | | 0.000 / NA | | | | |
| | AIC | 289.916 | | | | | 295.253 | | | | |

**Supplementary Table 12. Summary of a linear mixed-effects model testing whether later-life behavior predicts lifespan.** The model assesses the effects of activity and feeding behavior measured later in life, on individual lifespan.

| Response | Predictor | Full model | | | | | Final model | | | | |
|---|---|---|---|---|---|---|---|---|---|---|---|
| | | Estimate | SE | $\chi^2$ | p | df | Estimate | SE | $\chi^2$ | p | df |
| Lifespan (days) | (Intercept) | 2199.784 | 693.207 | - | - | - | 2229.995 | 694.060 | - | - | - |
| | Later-in-life activity (cm/sec, log-transformed) | 336.015 | 151.116 | 4.598 | 0.032 | 1 | 344.713 | 150.930 | 4.832 | 0.028 | 1 |
| | Squared later-in-life activity (cm/sec, log-transformed) | 18.014 | 8.109 | 4.590 | 0.032 | 1 | 18.483 | 8.098 | 4.826 | 0.028 | 1 |
| | Later-in-life feeding (min) | -0.290 | 0.539 | 0.291 | 0.591 | 1 | - | - | - | - | - |
| | **Random Effects** | | | | | | | | | | |
| | $\sigma^2$ | 5252.22 | | | | | 5299.84 | | | | |
| | $\tau_{00}$ | 0.00 Mother ID | | | | | 0.00 Mother ID | | | | |
| | *N* | 3 Mother ID | | | | | 3 Mother ID | | | | |
| | Observations | 32 | | | | | 32 | | | | |
| | Marginal $R^2$ / Conditional $R^2$ | 0.152 / NA | | | | | 0.144 / NA | | | | |
| | AIC | 356.623 | | | | | 355.601 | | | | |

**Supplementary Table 13. Linear mixed-effects model summaries testing whether early-life behavior predict lifetime reproductive output.** Separate models assessed effects of activity and feeding behavior on (i) reproductive onset, (ii) average offspring size across all broods, and (iii) average brood size across all broods produced over an individual's lifetime. Early-life activity and feeding were averaged across the first four weeks of life

| Response | Predictor | Full model | | | | | Final model | | | | |
|---|---|---|---|---|---|---|---|---|---|---|---|
| | | Estimate | SE | $\chi^2$ | p | df | Estimate | SE | $\chi^2$ | p | df |
| Reproductive onset (days) | (Intercept) | 135.221 | 14.686 | - | - | - | 132.994 | 8.060 | - | - | - |
| | Activity (cm/sec, log-transformed) | 2.767 | 7.470 | 0.137 | 0.714 | 1 | - | - | - | - | - |
| | Feeding (min) | -0.065 | 0.253 | 0.063 | 0.802 | 1 | - | - | - | - | - |
| | **Random Effects** | | | | | | | | | | |
| | $\sigma^2$ | 362.47 | | | | | 362.60 | | | | |
| | $\tau_{00}$ | 133.08 Mother ID | | | | | 152.33 Mother ID | | | | |
| | ICC | 0.27 | | | | | 0.30 | | | | |
| | *N* | 3 Mother ID | | | | | 3 Mother ID | | | | |
| | Observations | 33 | | | | | 33 | | | | |
| | Marginal $R^2$ / Conditional $R^2$ | 0.008 / 0.275 | | | | | 0.000 / 0.296 | | | | |
| | AIC | 291.900 | | | | | 293.005 | | | | |

**Supplementary Table 13.** Continued.

| Response | Predictor | Full model | | | | | Final model | | | | |
|---|---|---|---|---|---|---|---|---|---|---|---|
| | | Estimate | SE | $\chi^2$ | p | df | Estimate | SE | $\chi^2$ | p | df |
| Average brood size | (Intercept) | 15.893 | 3.780 | - | - | - | 15.707 | 1.741 | - | - | - |
| | Activity (cm/sec, log-transformed) | -2.149 | 1.939 | 1.082 | 0.298 | 1 | - | - | - | - | - |
| | Feeding (min) | 0.006 | 0.066 | 0.009 | 0.924 | 1 | - | - | - | - | - |
| | **Random Effects** | | | | | | | | | | |
| | $\sigma^2$ | 24.50 | | | | | 26.20 | | | | |
| | $\tau_{00}$ | 8.19 Mother ID | | | | | 6.15 Mother ID | | | | |
| | *N* | 0.25 | | | | | 0.19 | | | | |
| | Observations | 3 Mother ID | | | | | 3 Mother ID | | | | |
| | Marginal $R^2$ / Conditional $R^2$ | 33 | | | | | 33 | | | | |
| | AIC | 0.041 / 0.281 | | | | | 0.000 / 0.190 | | | | |
| | $\sigma^2$ | 210.940 | | | | | 208.088 | | | | |

**Supplementary Table 13.** Continued.

| Response | Predictor | Full model | | | | | Final model | | | | |
|---|---|---|---|---|---|---|---|---|---|---|---|
| | | Estimate | SE | $\chi^2$ | p | df | Estimate | SE | $\chi^2$ | p | df |
| Average offspring size (mm) | (Intercept) | 8.265 | 0.233 | - | - | - | 8.513 | 0.145 | - | - | - |
| | Activity (cm/sec, log-transformed) | 0.064 | 0.103 | 0.372 | 0.542 | 1 | - | - | - | - | - |
| | Feeding (min) | 0.005 | 0.003 | 1.995 | 0.158 | 1 | - | - | - | - | - |
| | **Random Effects** | | | | | | | | | | |
| | $\sigma^2$ | 0.07 | | | | | 0.07 | | | | |
| | $\tau_{00}$ | 0.07 Mother ID | | | | | 0.05 Mother ID | | | | |
| | *N* | 0.49 | | | | | 0.42 | | | | |
| | Observations | 3 Mother ID | | | | | 3 Mother ID | | | | |
| | Marginal $R^2$ / Conditional $R^2$ | 33 | | | | | 33 | | | | |
| | AIC | 0.037 / 0.513 | | | | | 0.000 / 0.425 | | | | |
| | $\sigma^2$ | 35.742 | | | | | 21.648 | | | | |

**Supplementary Table 14. Summary of a linear mixed-effects model testing whether lifetime reproductive output predicts lifespan.** Lifespan was modelled as a function of (i) age at reproductive onset, (ii) average offspring size across all broods, and (iii) average brood size across all broods produced over an individual's lifetime.

| Response | Predictor | Full model | | | | | Final model | | | | |
|---|---|---|---|---|---|---|---|---|---|---|---|
| | | Estimate | SE | $\chi^2$ | p | df | Estimate | SE | $\chi^2$ | p | df |
| Lifespan (days) | (Intercept) | 252.919 | 351.591 | - | - | - | 653.394 | 13.498 | - | - | - |
| | Reproductive onset (days) | 0.706 | 0.703 | 0.995 | 0.319 | 1 | - | - | - | - | - |
| | Average offspring size (mm) | 27.705 | 42.682 | 0.419 | 0.518 | 1 | - | - | - | - | - |
| | Average brood size | 4.792 | 2.536 | 3.390 | 0.066 | 1 | - | - | - | - | - |
| | **Random Effects** | | | | | | | | | | |
| | $\sigma^2$ | 5341.57 | | | | | 6012.78 | | | | |
| | $\tau_{00}$ | 0.00 Mother ID | | | | | 0.00 Mother ID | | | | |
| | *N* | 3 Mother ID | | | | | 3 Mother ID | | | | |
| | Observations | 33 | | | | | 33 | | | | |
| | Marginal $R^2$ / Conditional $R^2$ | 0.115 / NA | | | | | 0.000 / NA | | | | |
| | AIC | 368.015 | | | | | 379.746 | | | | |

#### Supplementary Note 8: Robustness analysis with respect to day of death uncertainty

**Supplementary Table 15. Robustness analysis excluding individuals with uncertain death dates.** Summary of linear mixed-effects models testing whether early-life activity and feeding behavior predict lifespan after excluding three individuals with uncertainty in death date. The model structure mirrors the one in the primary analysis (Supplementary Table 1). As in the primary analysis, behavior was first averaged across the entire four-week early-life observation period and then analyzed separately for each of the four weeks to assess the temporal development of the activity–lifespan relationship.

| **Time interval** | **Response** | **Predictor** | **Full model** | | | | | **Final model** | | | | |
|---|---|---|---|---|---|---|---|---|---|---|---|---|
| | | | **Estimate** | **SE** | **$\chi^2$** | **p** | **df** | **Estimate** | **SE** | **$\chi^2$** | **p** | **df** |
| Week 1-4 | Lifespan (days) | (Intercept) | 589.863 | 49.008 | - | - | - | 634.655 | 15.715 | - | - | - |
| | | Activity (cm/sec, log-transformed) | -140.496 | 40.812 | 9.988 | 0.002 | 1 | -151.627 | 39.743 | 11.866 | 0.001 | 1 |
| | | Squared Activity (cm/sec, log-transformed) | 141.277 | 46.941 | 7.916 | 0.005 | 1 | 140.887 | 47.659 | 7.669 | 0.006 | 1 |
| | | Feeding (min) | 0.870 | 0.904 | 0.914 | 0.339 | 1 | - | - | - | - | - |
| | | **Random Effects** | | | | | | | | | | |
| | | $\sigma^2$ | 4268.77 | | | | | 4400.81 | | | | |
| | | $\tau_{00}$ | 0.00 Mother ID | | | | | 0.00 Mother ID | | | | |
| | | *N* | 3 Mother ID | | | | | 3 Mother ID | | | | |
| | | Observations | 30 | | | | | 30 | | | | |
| | | Marginal $R^2$ / Conditional $R^2$ | 0.356 / NA | | | | | 0.336 / NA | | | | |
| | | AIC | 321.603 | | | | | 322.185 | | | | |

**Supplementary Table 15.** Continued.

| Time interval | Response | Predictor | Full model | | | | | Final model | | | | |
|---|---|---|---|---|---|---|---|---|---|---|---|---|
| | | | Estimate | SE | $\chi^2$ | p | df | Estimate | SE | $\chi^2$ | p | df |
| Week 1 | Lifespan (days) | (Intercept) | 664.945 | 38.039 | - | - | - | 699.022 | 18.920 | - | - | - |
| | | Activity (cm/sec, log-transformed) | -75.271 | 66.122 | 1.269 | 0.260 | 1 | -65.905 | 19.576 | 9.614 | 0.002 | 1 |
| | | Squared Activity (cm/sec, log-transformed) | 16.838 | 39.500 | 0.181 | 0.670 | 1 | - | - | - | - | - |
| | | Feeding (min) | 0.932 | 0.807 | 0.305 | 0.253 | 1 | - | - | - | - | - |
| | | **Random Effects** | | | | | | | | | | |
| | | $\sigma^2$ | 4502.72 | | | | | 4753.21 | | | | |
| | | $\tau_{00}$ | 0.00 Mother ID | | | | | 0.00 Mother ID | | | | |
| | | *N* | 3 Mother ID | | | | | 3 Mother ID | | | | |
| | | Observations | 30 | | | | | 30 | | | | |
| | | Marginal $R^2$ / Conditional $R^2$ | 0.320 / NA | | | | | 0.281 / NA | | | | |
| | | AIC | 324.065 | | | | | 332.376 | | | | |

**Supplementary Table 15.** Continued.

| Time interval | Response | Predictor | Full model | | | | | Final model | | | | |
|---|---|---|---|---|---|---|---|---|---|---|---|---|
| | | | Estimate | SE | $\chi^2$ | p | df | Estimate | SE | $\chi^2$ | p | df |
| Week 2 | Lifespan (days) | (Intercept) | 651.109 | 36.087 | - | - | - | 649.199 | 13.741 | - | - | - |
| | | Activity (cm/sec, log-transformed) | -98.854 | 32.563 | 8.037 | 0.005 | 1 | -52.730 | 23.866 | 4.523 | 0.033 | 1 |
| | | Squared Activity (cm/sec, log-transformed) | 63.820 | 34.456 | 3.248 | 0.072 | 1 | - | - | - | - | - |
| | | Feeding (min) | -0.466 | 0.603 | 0.591 | 0.442 | 1 | - | - | - | - | - |
| | | **Random Effects** | | | | | | | | | | |
| | | $\sigma^2$ | 4983.97 | | | | | 5632.44 | | | | |
| | | $\tau_{00}$ | 0.00 Mother ID | | | | | 0.00 Mother ID | | | | |
| | | *N* | 3 Mother ID | | | | | 3 Mother ID | | | | |
| | | Observations | 30 | | | | | 30 | | | | |
| | | Marginal $R^2$ / Conditional $R^2$ | 0.245 / NA | | | | | 0.144 / NA | | | | |
| | | AIC | 327.524 | | | | | 336.901 | | | | |

**Supplementary Table 15.** Continued.

| Time interval | Response | Predictor | Full model | | | | | Final model | | | | |
|---|---|---|---|---|---|---|---|---|---|---|---|---|
| | | | Estimate | SE | $\chi^2$ | p | df | Estimate | SE | $\chi^2$ | p | df |
| Week 3 | Lifespan (days) | (Intercept) | 491.471 | 55.049 | - | - | - | 572.286 | 30.938 | - | - | - |
| | | Activity (cm/sec, log-transformed) | -89.828 | 36.797 | -2.441 | 0.022 | 1 | -93.445 | 38.544 | -2.424 | 0.023 | 1 |
| | | Squared Activity (cm/sec, log-transformed) | 136.658 | 45.609 | 2.996 | 0.006 | 1 | 136.477 | 47.851 | 2.852 | 0.009 | 1 |
| | | Feeding (min) | 1.302 | 0.749 | 1.739 | 0.095 | 1 | - | - | - | - | - |
| | | **Random Effects** | | | | | | | | | | |
| | | $\sigma^2$ | 4679.87 | | | | | 5151.38 | | | | |
| | | $\tau_{00}$ | 0.00 Mother ID | | | | | 0.00 Mother ID | | | | |
| | | *N* | 3 Mother ID | | | | | 3 Mother ID | | | | |
| | | Observations | 30 | | | | | 30 | | | | |
| | | Marginal $R^2$ / Conditional $R^2$ | 0.292 / NA | | | | | 0.219 / NA | | | | |
| | | AIC | 324.926 | | | | | 326.902 | | | | |

**Supplementary Table 15.** Continued.

| Time interval | Response | Predictor | Full model | | | | | Final model | | | | |
|---|---|---|---|---|---|---|---|---|---|---|---|---|
| | | | Estimate | SE | $\chi^2$ | p | df | Estimate | SE | $\chi^2$ | p | df |
| Week 4 | Lifespan (days) | (Intercept) | 613.985 | 41.202 | - | - | - | 651.467 | 14.775 | - | - | - |
| | | Activity (cm/sec, log-transformed) | 9.480 | 41.106 | 0.053 | 0.818 | 1 | - | - | - | - | - |
| | | Squared Activity (cm/sec, log-transformed) | 19.067 | 54.699 | 0.121 | 0.728 | 1 | - | - | - | - | - |
| | | Feeding (min) | 0.637 | 0.758 | 0.698 | 0.404 | 1 | - | - | - | - | - |
| | | **Random Effects** | | | | | | | | | | |
| | | $\sigma^2$ | 6283.47 | | | | | 6548.92 | | | | |
| | | $\tau_{00}$ | 0.00 Mother ID | | | | | 0.00 Mother ID | | | | |
| | | *N* | 3 Mother ID | | | | | 3 Mother ID | | | | |
| | | Observations | 30 | | | | | 30 | | | | |
| | | Marginal $R^2$ / Conditional $R^2$ | 0.042 / NA | | | | | 0.000 / NA | | | | |
| | | AIC | 332.559 | | | | | 347.507 | | | | |

### Supplementary Note 9: Protocol of standard behavioral assays

Immediately following the 28-day period of continuous behavioral observations, individuals underwent a series of standardized behavioral assays conducted once per day (starting on day 29). Each day, a single assay was performed—activity in a novel tank, sociability, or response to a novel object—using the same automated, high-resolution recording system employed during the initial observation phase. The assays were conducted in the following order and then repeated two more times in the same order (i.e., any particular test was conducted a total of three times). On day 1, individuals were assayed for their activity in a new tank. On day 2, individuals experienced either a sociability test or a novel object test in randomized order. On day 3, individuals experienced whichever of the sociability or novel object tests was not experienced on day 2.

To initiate the new tank activity assay, all tanks were cleaned and individuals were swapped among observation tanks. Activity (average distance moved per second, in cm) was tracked over a period of 60 minutes, starting one minute after transfer. For the following sociability and novel object test, individuals remained in the same tank used for the new tank activity assay. Care was taken to ensure that individuals experienced each of a total of four tank systems (three systems in addition to the tank system in which they were kept for the first 28 days) as well as all possible tank positions (center vs. periphery). Due to technical issues, we excluded $N$ = 5 trials, i.e., $N$ (final) = 97 new tank trials.

Sociability tests were conducted by introducing a round, clear acrylic container (10 cm diameter, 10 cm tall) containing two conspecifics into the tank. Conspecifics were matched to the size of stimulus fish as closely as possible (mean ± SD total length = 1.8 ± 0.3 cm; $N$ = 80 randomly chosen from $N$ = 176 stimulus fish used) but were otherwise randomly chosen from a stock tank. Each conspecific stimulus fish was used only once during the course of the experiment. Additionally, in order to control for a potential effect of the container itself, an empty container was also introduced to the tank, with the position of the empty container vs. the container containing conspecifics being randomized between two possible positions. The focal individual and the stimulus fish were given a 10-minute acclimation period, during which containers were surrounded with opaque white flower pots, followed by a 60-minute trial period (starting one minute after the flower pots were gently removed). Sociability was assessed as an individual's relative preference for conspecifics over the empty container. Specifically, sociability was quantified as the proportion of time spent within a 5-cm distance to the conspecific container, relative to the total time spent within a 5-cm distance to either container. For $N$ = 7 trials, we could not calculate individual relative preference as focal fish visited neither container; and due to technical issues, we excluded $N$ = 7 trials, i.e., $N$ (final) = 88 sociability trials.

Novel object tests were conducted by introducing one of the following three novel objects to the tanks (such that every fish experienced each of the three objects once over the course of three novel object assays in a randomized order): a clear bounce ball filled with star-shaped colorful glitter specks (red, green, yellow, blue; ball diameter = 4.3 cm); a ceramic, light green flower pot (diameter = 8 cm, height = 7 cm), and a plastic pink anemone (diameter = 10 cm). Individual responses to these objects were assessed as the average distance (cm) to the object over 60 minutes (starting 1 minute after introducing the object). Due to technical issues, we excluded $N$ = 2 trials, i.e., $N$ (final) = 100 novel object trials.
